# Post-fibrillization nitration of alpha-synuclein abolishes its seeding activity and pathology formation in primary neurons and *in vivo*

**DOI:** 10.1101/2023.03.24.534149

**Authors:** Sonia Donzelli, Sinead A. OSullivan, Anne-Laure Mahul-Mellier, Ayse Ulusoy, Giuliana Fusco, Senthil T. Kumar, Anass Chiki, Johannes Burtscher, Manel L.D. Boussouf, Iman Rostami, Alfonso De Simone, Donato A. Di Monte, Hilal A. Lashuel

## Abstract

Increasing evidence points to post-translational modifications (PTMs) as key regulators of alpha-synuclein (α-Syn) function in health and disease. However, whether these PTMs occur before or after α-Syn pathology formation and their role in regulating α-Syn toxicity remain unclear. In this study, we demonstrate that post-fibrillization nitration of α-Syn fibrils induced their fragmentation, modified their surface and dynamic properties but not their structure, and nearly abolished their seeding activity in primary neurons and *in vivo*. Furthermore, we show that the dynamic and surface properties of the fibrils, rather than simply their length, are important determinants of α-Syn fibril seeding activity. Altogether, our work demonstrates that post-aggregation modifications of α-Syn may provide novel approaches to target a central process that contributes to pathology formation and disease progression. Finally, our results suggest that the pattern of PTMs on pathological aggregates, rather than simply their presence, could be a key determinant of their toxicity and neurodegeneration. This calls for reconsidering current approaches relying solely on quantifying and correlating the level of pathology to assess the efficacy of novel therapies, as not all α-Syn aggregates in the brain are pathogenic.

## Introduction

Protein tyrosine nitration and dityrosine crosslinking are oxidative and nitrosative stress biomarkers which have been implicated in normal aging and the pathogenesis of numerous neurodegenerative disorders (NDDs)^1–4^. In these diseases, the production of reactive oxygen (ROS) and nitrogen (RNS) species overwhelms the antioxidant capacity of cells^5^, leading to a redox imbalance and ultimately to increased production of the nitrogen dioxide radical (NO_2_), 3-nitrotyrosine, and dityrosine cross-linking formation^3, 6^. Several neuropathological observations and recent rodent studies also suggest this to be the case in Parkinson’s disease (PD)^7–9^, the second most common age-related NDD.

PD is characterized by the selective vulnerability of specific neuronal populations, particularly dopaminergic neurons in the substantia nigra pars compacta (SNpc), where high levels of ROS and RNS have been proposed to contribute to the loss of these neurons^10^. One of the main pathological hallmarks of PD is the presence of Lewy bodies (LBs) and Lewy neurites, intraneuronal inclusions that are enriched in aggregated and post-translationally modified forms of the protein α-synuclein (α-Syn)^11, 12^. Widespread accumulation of 3-nitrotyrosine and dityrosine cross-linked α-Syn aggregates and fibrils has been observed in the brains of patients with PD^2, 4, 13–16^, dementia with LBs (DLBs)^17^, and multiple system atrophy (MSA)^2, 18, 19^, suggesting an association between oxidative and nitrative stress, and α-Syn aggregation and pathology formation in PD and related synucleinopathies. Furthermore, a 9-fold increase in α-Syn nitration at Y39 has been reported upon overexpression of monoamine oxidase B in animal models of PD^20^. In several cellular and genetic and toxin-induced models of synucleinopathies, increasing oxidative stress reportedly regulates α-Syn toxicity and cell-to-cell-transmission^21, 22^, and/or enhances α-Syn aggregation or inclusion formation^23, 24^. Increased accumulation of nitrated α-Syn within dopaminergic neurons has also been reported to occur during normal aging in the primate brain, but whether this reflects toxic or neuroprotective mechanisms remains unclear^25^.

The pervasive presence of nitrated α-Syn in PD patients’ brains^2, 4, 18^ led to the initial hypothesis that α-Syn nitration occurs at the onset of α-Syn aggregation and may contribute to the formation of LBs and/or PD pathogenesis. Therefore, initial studies aimed at exploring the links between these processes focused primarily on investigating the effects of tyrosine nitration and dityrosine cross-linking on the aggregation and fibrillization of α-Syn monomers *in vitro*. These studies showed that the nitration of monomeric α-Syn results in the intermolecular dityrosine-mediated formation of partially folded states and stable oligomeric species that do not go on to form α-Syn fibrils^26–30^. In these studies, non-specific chemical nitration of all tyrosine residues of α-Syn resulted in the generation of heterogeneous mixtures of α-Syn species with different nitration patterns, thus making it challenging to dissect the role of nitration at each tyrosine residue. To address these limitations, we previously developed semisynthetic protein strategies that enabled the site-specific nitration of α-Syn^30^. *In vitro* aggregation studies showed that α-Syn nitrated at Y39 (nY39) or Y125 (nY125) formed β-sheet rich fibrils with similar aggregation kinetics as the wild-type (WT) unmodified protein. However, the fibrils formed by the Y39 or Y125 nitrated proteins exhibited distinct structures and morphological features compared to those formed by the unmodified protein or the protein extracted post-mortem from disease-affected brains^30^. Both nY39 and nY125 α-Syn formed predominantly short and thick fibrils, particularly in the case of nY39, whereas nY125 aggregates exhibited a high propensity to stack in parallel. Taken together, these findings suggest that the non-specific nitration of monomeric α-Syn interferes with its fibrillization and stabilizes oligomeric forms of the protein. By contrast, site-specific nitration at the N-or C-terminus significantly alters the final structure and morphology of the fibrils. This latter effect suggests that site-specific nitration could play an important role in determining the final α-Syn fibril strain.

Nitration of α-Syn in pathological aggregates (LBs and LNs) could also occur after α-Syn fibrillization and/or LB formation. Indeed, increasing evidence suggests that several α-Syn C-terminal post-translational modifications (PTMs) found in LB inclusions (e.g., ubiquitination, phosphorylation, and truncation) are not required for α-Syn aggregation and occur after fibril formation and/or during the transition from fibrils to LBs^31, 32^. However, the effects of post-fibrillization PTMs, including nitration, on the structure, stability, seeding capacity, and remodeling of fibrils have not been investigated. In this work, we sought to address this knowledge gap by conducting a systematic investigation of the effects of post-fibrillization nitration on the structural properties, morphology, and fragmentation of α-Syn fibrils and on α-Syn fibril seeding and spreading activity *in vitro* and *in vivo*. To the best of our knowledge, this is the first study to address the role of post-fibrillization nitration on these aspects of α-Syn aggregation and pathology formation. Our findings reveal that post-aggregation nitration of α-Syn pre-formed fibrils (PFFs) induces their fragmentation and neutralizes their seeding activity in primary neurons and *in vivo* without altering their structure. These findings point to the important roles of PTMs in regulating α-Syn pathology formation and spreading in PD and provide intriguing new evidence supporting a neuroprotective effect of post-fibrillation nitration during pathogenetic processes in human synucleinopathies.

## Results

### Post-fibrillization nitration of α-Syn fibrils does not alter their structure or morphology

To determine if post-aggregation nitration influences the seeding activity of α-Syn fibrils, we first generated α-Syn PFFs from recombinant monomeric human α-Syn as described in Kumar et al.^33^ (Fig. 1A–C). Half of the fibril preparations were subjected to *in vitro* nitration using tetranitromethane (TNM) (Fig. 1D–F). The formation of 3-nitrotyrosine, as a result of tyrosine nitration, was monitored by electrospray ionization mass spectrometry (ESI-MS) (Fig. 1D), and the human nitrated (Ni) and unmodified WT α-Syn were characterized by western blotting (WB) (Fig. 1E and Supplemental Fig. 1) and transmission electron microscopy (TEM) analyses (Fig. 1F). ESI-MS analysis of human WT α-Syn prior to nitrative modification yielded one major peak (14,458 Da) corresponding to unmodified α-Syn. TNM-mediated nitration of α-Syn resulted in two α-Syn fibril populations with either three or four tyrosine residues being nitrated, as evidenced by the two major MS peaks positioned at 14,593 Da (14,458 + 135 [3 NO_2_]) and 14,638 Da (14,458 + 180 [4 NO_2_]), respectively. Since three of the tyrosine residues of α-Syn (Y125/Y133/Y136) are in the C-terminal domain, which remains flexible and accessible in the fibrillar state, we suspected that the tri-nitrated species corresponded to α-Syn fibrils with all C-terminal tyrosine residues nitrated. The fourth tyrosine residue, Y39, which is in close proximity to the core of the fibrils, is less accessible and thus may require longer incubation times to achieve complete nitration. Therefore, the ESI-MS results indicated that the C-and N-terminal tyrosine residues in α-Syn may not be equally susceptible to nitration once α-Syn fibrils or LBs are formed. Our results are in line with the location of these residues in several of the reported cryogenic electron microscopy (cryo-EM) structures of recombinant α-Syn fibrils, which exhibit a core sequence spanning residues 37–99 and a highly flexible C-terminal domain (residues 100–140)^34–36^. Furthermore, these studies are also consistent with previous nuclear magnetic resonance (NMR) studies reporting that the C-terminal region of α-Syn fibrils surrounds the structured core of the fibril with a dense mesh of disordered tails^37^.

**Figure 1:**
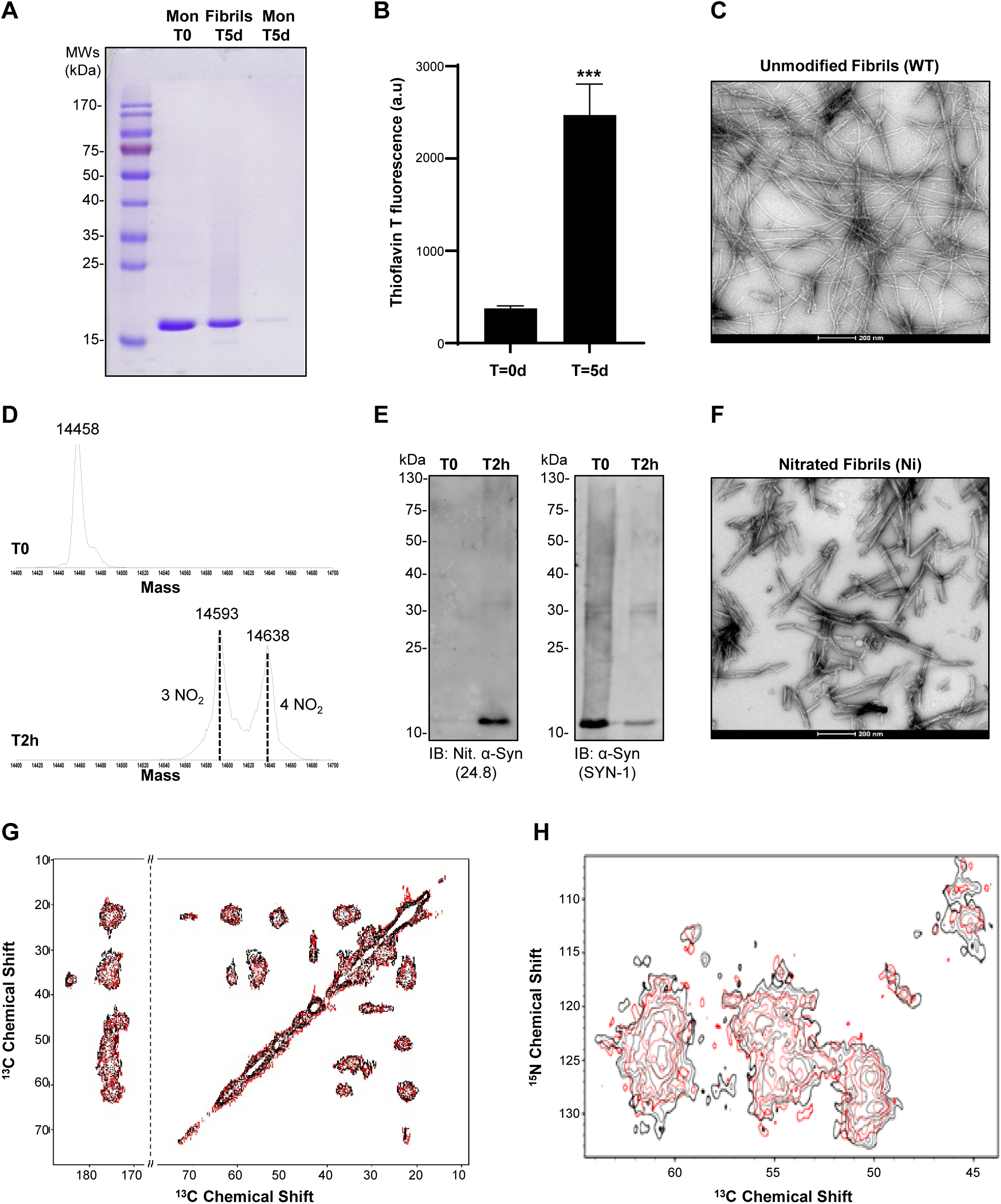
*In vitro* nitration did not alter the conformational properties of α-Syn PFFs. Generation of α-Syn fibrils from recombinant α-Syn. **A.** Coomassie staining of monomeric α-Syn (first line), mature fibrils after 5 days of aggregation (second line), and monomers that did not aggregate (third line). **B.** ThT fluorescence of monomeric α-Syn (T = day 0) and mature fibrils (T = day 5). **C.** TEM of mature fibrils before nitration. **D.** ESI-LC/MS analysis of the extent of TNM-mediated nitration of α-Syn fibrils after 2 h of incubation. α-Syn fibrils were boiled at 95°C for 10 min and centrifuged (max speed, 4°C, 10 min), after which the supernatant was subjected to ESI-MS for analysis. **E.** WB analysis of unmodified (T = 0) and nitrated fibrils (T2h). Antibody 24.8 was used to detect postfibrillization nitration on fibrils and SYN-1 was used to reveal the total amount of α-Syn. **F.** TEM of mature fibrils after nitration. **G.** ssNMR spectra of unmodified (red) and nitrated (black) α-Syn fibrils. ^13^C-^13^C-DARR spectra were recorded with a 50 ms mixing time at 278 K. These experiments are accurate probes of the structure of core regions of amyloid fibrils. The line shapes of the spectra indicate that the fibrils possess a level of local conformational heterogeneity. This is consistent with previous ssNMR studies of α-Syn fibrils made in the absence of seeding. The overlap of the spectra revealed a complete match between the resonances of the two fibrils, indicating that the sequences and conformations of the core regions of the two types of fibrils matched. **H.** ^13^C-^15^N NCA ssNMR spectra of unmodified (red) and nitrated (black) α-Syn fibrils studied in this work. The two spectra showed a significant overlap of the resonances, in agreement with the ^13^C–^13^C-DARR spectra.

Given that the majority of tyrosine residues on the fibrils become nitrated, we sought to determine whether this could lead to the remodeling of fibril morphology and structure over time. To test this hypothesis, we 1) assessed changes in the morphology and width of nitrated fibrils over time; 2) compared the proteinase K (PK) resistance profile of nitrated fibrils; 3) and performed solid-state NMR (ssNMR) to compare the structure of the same α-Syn PFF before and after nitration. As shown in Supplemental Figure 2, we did not observe any significant changes in fibril morphology (Supplemental Fig. 2A) upon nitration of fibrils derived from WT proteins (Supplemental Fig. 2C). Although both unmodified and nitrated PFFs exhibited similar sodium dodecyl-sulfate polyacrylamide gel electrophoresis (SDS-PAGE) profiles, the nitrated fibrils showed bands with reduced intensity in addition to SDS and PK-resistant high molecular weight (HMW) bands (Supplemental Fig. 2B), most likely due to nitration-induced cross-linking of the fibrils. Interestingly, the nitration of PFFs resulted in an immediate reduction in their length and width that persisted up to 35 days (Supplemental Fig. 2B). These observations prompted us to further characterize and compare the structural properties of ^15^N^13^C WT and nitrated ^15^N^13^C WT α-Syn fibrils (Supplemental Fig. 3A–E) using ssNMR coupled with magic angle spinning (MAS) experiments. ^13^C-^13^C dipolar-assisted rotational resonance (DARR) correlation spectra were measured to probe highly rigid regions of the fibril cores. The resulting distribution of chemical shifts in the spectra is consistent with previous ssNMR analyses of α-Syn fibrils, which revealed a characteristic Greek key structural topology^38^. The significant overlap of the ^13^C-^13^C DARR spectra of WT and nitrated α-Syn fibrils indicated a strong match in the structural properties of these two types of fibrils (Fig. 1G). Individual spectra are shown in Supplemental Figure 3F. Similar resonance matches between the WT and nitrated α-Syn fibrils spectra were found for heteronuclear NCA spectra (Fig. 1H), with individual spectra shown in Supplemental Figure 3G. Taken together, these ssNMR spectra indicate that the sequence and conformations of the core regions of the nitrated α-Syn fibrils are virtually identical to those of the unmodified WT fibrils.

### Post-fibrillization nitration of α-Syn PFFs significantly reduces their seeding activity, propagation, and ability to induce neurodegeneration in mouse brains

Potential differences in the seeding and pathological features between unmodified and nitrated α-Syn fibrils were first investigated *in vivo* in mice treated with a unilateral intraparenchymal injection of either unmodified or nitrated human PFFs (batch #1; Supplemental Fig. 4) into the right striatum. Animals were sacrificed at 12 or 20 weeks post-injection (Fig. 2A). Immunostaining with an antibody that specifically recognizes α-Syn phosphorylated at Ser129 (pS129-α-Syn) was used as a marker of pathological accumulation of the protein^39–42^. Coronal sections of the striatum from unmodified PFF-injected mice revealed overt pathological features in the form of perikaryal and neuritic p129-α-Syn-labeled inclusions (Fig. 2B). pS129-α-Syn-positive cell bodies appeared to be most densely clustered in proximity to the injection site. Microscopic analysis of specimens from animals treated with nitrated α-Syn PFFs showed strikingly different results, with very little pathology observed after the staining of striatal tissue sections with anti-pS129-α-Syn (Fig. 2B). Semi-quantitative densitometric measurements of pS129-α-Syn immunoreactivity confirmed the different effects of unmodified versus nitrated PFFs; the former induced robust pS129-α-Syn pathology at both the 12-and 20-week time points, whereas very little immunoreactivity characterized the striatal tissue after injection of nitrated PFFs (Fig. 2C). The amygdala and substantia nigra are preferential targets of α-Syn pathology in PD, and based on the findings of earlier investigations, are also highly vulnerable to pS129-α-Syn accumulation after striatal PFFs administration^39, 43^. Therefore, we conducted further analyses of pathological α-Syn accumulation in coronal sections of the amygdala and substantia nigra from PFFs-injected mice (Fig. 2D–G). When these sections were stained with anti-pS129-α-Syn, robust immunoreactivity was seen within both neuritic processes and neuronal cell bodies in all samples from unmodified PFFs-injected animals (Fig. 2D, F). pS129-α-Syn-positive inclusions varied in size, shape, and intensity; for example, some neurons displayed relatively large, compact, and crescent-shaped inclusions, whereas other cells were characterized by grainier immunoreactivity (Supplemental Fig. 5). Results in the amygdala and substantia nigra of mice treated with nitrated α-Syn PFFs were unequivocally different since both microscopic observations and semi-quantitative measurements detected only minimal pS129-α-Syn pathology (Fig. 2D–G).

**Figure 2:**
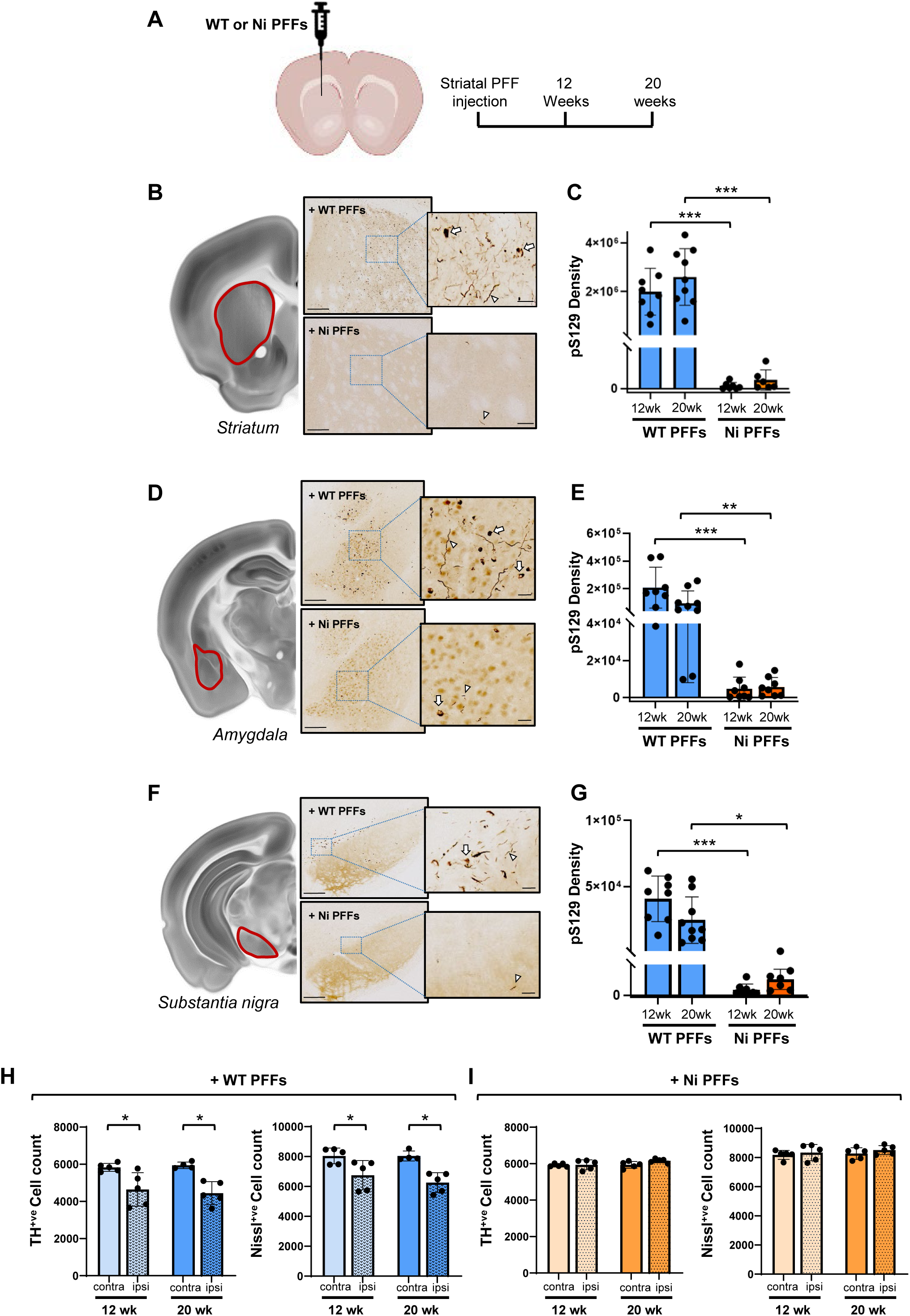
Reduced seeding capability of nitrated compared to unmodified PFFs *in vivo*. **A.** Experimental study design. Unmodified and nitrated human PFFs (5 mg) were injected unilaterally into the mouse striatum, and animals were sacrificed at 12 or 20 weeks post-injection. **B, D and F.** Schematic representations of the areas of interest are depicted and delineated in red: striatum (**B**), amygdala (**D**) and substantia nigra (**F**). Tissue sections from these regions of interest (ipsilateral to the PFF injection side) were collected at 12 weeks and processed for histology. Representative microscopy images at low and high magnification show brain sections stained with an anti-pS129 α-syn antibody (EP1536Y). Examples of perikaryal (arrows) and neuritic pathology (arrowheads) are shown within the high-magnification images. Scale bars: 200 μm (low magnification images) and 20 μm (higher magnification images). **C, E and G.** pS129-α-Syn pathology was quantified according to pixel density in sections from the striatum (**C**), amygdala (**E**) and substantia nigra (**G**) of mice injected with either unmodified or nitrated PFFs (n=8-9 animals/experimental group). Columns indicate means, and error bars indicate SD. ***P < 0.001, **P < 0.002. (**H and I**) Midbrain tissue sections containing the substantia nigra were collected from mice injected with either unmodified or nitrated PFFs (n = 5 animals/experimental group) and stained with anti-tyrosine hydroxylase (TH) and cresyl violet. Then TH-positive neurons and Nissl-stained cells were counted in the substantia nigra pars compacta using unbiased stereological techniques. Counts in the substantia nigra ipsilateral and contralateral to the PFF injection site are shown. Columns indicate means, and error bars indicate standard deviation (SD). *P < 0.03.

In both PD and rodent PD models, dopaminergic neurons in the SNpc are not only susceptible to the formation of pathological α-Syn inclusions but are also highly vulnerable to neurodegenerative processes^44, 45^. Therefore, we assessed the effects of striatal injections of either unmodified or nitrated α-Syn PFFs on the integrity/survival of nigral dopaminergic cells. The number of these cells was counted in midbrain tissue sections spanning the entire SNpc using a stereological approach. The sections were stained with an antibody against tyrosine hydroxylase (TH, a marker of dopaminergic cells) and counterstained with cresyl violet. The number of TH-positive or Nissl-stained cells was significantly decreased by 20–25% in the right (PFF-injected side) SNpc from unmodified PFFs-treated mice at the two analyzed time points (Fig. 2H). By contrast, the number of nigral neurons remained unchanged after the administration of nitrated PFFs (Fig. 2I). Taken together, these findings demonstrate that PTMs of fibrillar α-Syn markedly modulate its pathological effects *in vivo*. In particular, the ability of PFFs to induce α-Syn propagation, intra-neuronal protein deposition, and neuronal injury was markedly attenuated as a consequence of nitrative modifications of the fibrillar protein.

### Post-fibrillization nitration does not influence the uptake, processing or clearance of α-Syn PFFs

Having established that the seeding capacity of human PFFs is dramatically reduced by their nitration, we sought to gain insights into the specific mechanisms by which the nitration of α-Syn PFFs could affect the induction and spreading of α-Syn pathology. We investigated whether post-fibrillization nitration modulates PFFs uptake, proteolytic processing, and the ability of PFFs to promote α-Syn fibrillization and the formation of LB-like inclusions. For these experiments, we used a cellular model of neuronal seeding, namely primary neuronal cultures in which exogenous α-Syn PFFs induce intraneuronal aggregation of endogenous α-Syn^46, 47^. The seeding process is initiated only after the internalization of PFFs into neurons, which consequently leads to the formation of predominantly fibrillar α-Syn aggregates (between Day 7 [D7] and D14). These aggregates eventually (on D21) form LB-like inclusions that exhibit morphological, biochemical, and ultrastructural features of LBs in PD brains^46, 48^. First, we investigated whether nitration influences the uptake and proteolytic processing of PFFs. The use of α-Syn knock-out (KO) mouse-derived primary neurons allowed us to follow the fate of human α-Syn PFFs independently of the seeding process and formation of new aggregates due to the absence of endogenous α-Syn expression. Immunocytochemistry (ICC) combined with confocal microscopy analysis confirmed that Ni and WT α-Syn PFFs (batch #2; Supplemental Fig. 4) were both internalized into microtubule-associated protein 2 (MAP2)-positive neurons through the endolysosomal pathway as previously described^46^ (Fig. 3A). Then insoluble fractions of the KO neurons were analyzed by WB for up to 10 days. Figure 3B confirms the ICC data and shows that within 24 h, both the nitrated and unmodified PFFs were readily internalized. WB analysis further revealed SDS-resistant aggregates of a larger size (HMW species) that were more readily detected by a total α-Syn antibody (4B12) in the insoluble fraction of KO neurons treated with Ni PFFs than those treated with WT PFFs (Fig. 3B, upper panel and Supplemental Fig. S6). The significantly reduced levels of monomeric α-Syn (15 kDa) in nitrated PFFs-treated neurons (Fig. 3B, middle panel) is expected given the fact that they undergo substantial intra-and inter-molecular cross-linking, as evidenced by the significantly higher accumulation of SDS-resistant HMW aggregates (Fig. 3B, upper panel and Supplemental Fig. S6) as previously reported^30^.

**Figure 3:**
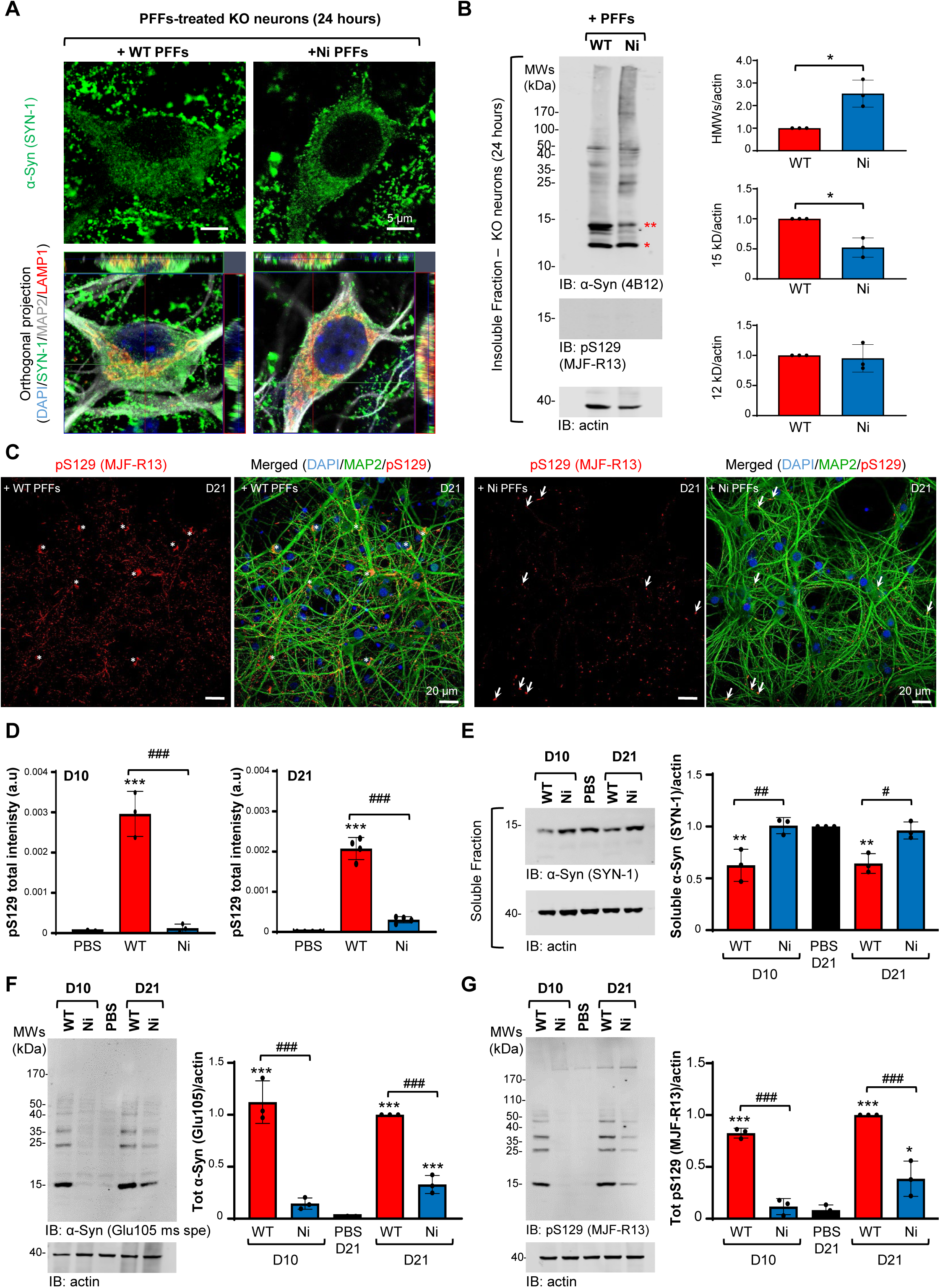
Uptake, processing, and seeding capacity of the WT and nitrated PFFs in primary hippocampal neurons. α-Syn KO (**A-B**) or WT (**C-G**) hippocampal primary neurons were treated with 70 nM of WT or Ni α-Syn PFFs for 24 h (D1, **A-B**) and up to 21 days (D21, **C-G**). Control neurons were treated with PBS. The uptake, processing, and seeding capacity of the PFFs were evaluated by ICC (**A, C**), HCA (**D**) and WB (**B, E–G**) analyses. **A.** ICC analysis show the internalization of WT and Ni α-Syn PFFs into the MAP2-positive α-Syn KO neurons. The PFFs were detected by total α-Syn (SYN-1) antibody, the neurons were counterstained with MAP2 antibody, and the nuclei with DAPI. The late endosome/lysosomes were stained with LAMP1 antibody. Scale bars = 5 μm. **B.** WB analysis of the insoluble fractions of α-Syn KO neurons treated with WT and Ni α-Syn PFFs for 24 h showed the detection of the 12 kDa (one red star), 15 kDa (double red stars), and HMWs bands by the total α-Syn antibody (4B12). The levels of α-Syn were normalized to actin. The densitometry analysis is displayed on the right-hand side. **C.** The seeded aggregates formed in the cell bodies (white star) and the neurites (white arrows) of the MAP2-positive neurons were detected by ICC using the pS129 antibody (MJF-R13) and imaged by confocal imaging. Scale bars = 20 μm. **D.** The level of pS129-α-Syn-positive seeded aggregates was quantified at D10 and D21 in PBS-and PFFs-treated WT neurons by HCA. The seeded aggregates were detected by ICC using pS129 (81A) antibody, the neurons were counterstained with MAP2 antibody, and the nuclei with DAPI (see representative images in Supplemental Fig. 7). For each independent experiment (n = 3), a minimum of two wells was acquired per condition, and nine fields of view were imaged per well. **E.** WB analysis of the soluble fractions of WT neurons treated with WT or Ni PFFs showed a significant decrease of the endogenous α-Syn (15 kDa), specifically in the WT PFF-treated neurons. The levels of α-Syn were normalized to the actin. **F-G.** WB analysis of the insoluble fractions of WT neurons treated with WT or Ni human PFFs. The level of seeded aggregates was detected with an antibody specific for mouse α-Syn (Glu105) (**F**) or with a pS129 antibody (MJF-R13) (**G**). The levels of α-Syn or pS129 α-Syn were normalized to the actin. **B, E-G.** The graphs represent the mean +/- SD of a minimum of three independent experiments. **B.** *P < 0.01 (two-tailed paired t-test). **C, E-G.** *P < 0.01, **P < 0.001, ***P < 0.0001 (one-way ANOVA followed by Tukey’s HSD posthoc test, PBS vs. PFF-treated neurons). ### P < 0.0001 (one-way ANOVA followed by Tukey’s HSD posthoc test, WT PFF-treated neurons vs. Ni PFF-treated neurons).

Consistent with the results of previous studies^32, 46, 49^, the internalized WT and Ni PFFs were both efficiently cleaved at the C-terminus 24 h post-treatment, leading to the accumulation of 12 kDa fragments (indicated by a red star in Fig. 3B and Supplemental Fig. S6). After 3 days, full-length α-Syn (15 kDa) levels were strongly reduced in both WT and Ni PFFs-treated neurons; this reduction was accompanied by a marked increase in cleaved protein (12 kDa) only in cultures treated with WT PFFs (Supplemental Fig. S6). Truncated species (12 kDa) for both types of PFFs were still detected in the KO neurons 10 days after PFFs treatment, while levels of full-length α-Syn (15 kDa) continued to decrease and completely disappeared between D7 and D10 in both conditions (Supplemental Fig. S6). Finally, and as previously described ^32^, the internalized PFFs were never phosphorylated (at residue S129) in KO neurons, even at early time points (i.e., prior to cleavage of the C-terminal domain) (Supplemental Fig. S6). Together, these data demonstrate that post-fibrillization nitration does not interfere with the uptake of α-Syn PFFs, their processing (C-terminal cleavage), or the general patterns of their clearance after internalization.

### Post-fibrillization nitration reduces the seeding capacity of α-Syn PFFs in primary neurons

Next, we evaluated and compared the seeding capacity of WT and nitrated PFFs using ICC combined with high content imaging analysis (HCA) in primary cultures of hippocampal neurons from WT mice. This approach allows the quantification of pS129-α-Syn-positive newly formed aggregates in MAP2-positive neurons (Fig. 3C and Supplemental Fig. S7A). In sharp contrast to WT PFFs, the nitrated PFFs barely induced any seeding of newly formed aggregates at 10 days after their addition to WT neurons (Fig. 3C and Supplemental Fig. S7A). Indeed, while pS129-positive aggregates were detected in both neurites and cell bodies of WT PFFs-treated cultures, only a few seeded-induced pS129-α-Syn-positive aggregates were formed within Ni PFFs-treated neurons; these Ni PFFs-induced aggregates were mostly confined to neurites (Fig. 3C and Supplemental Fig. S7A). Even 21 days after the addition of nitrated PFFs, the formation of pS129-α-Syn-positive seeded aggregates remained significantly lower than levels observed in WT PFFs-treated neurons (Fig. 3C and Supplemental Fig. S7A), as quantified in Figure 3D.

To further validate our ICC data, we monitored the seeding activity of WT and Ni PFFs on D10 and D21 by WB analyses (Fig. 3E–G and Supplemental Fig. S7B). First, levels of endogenous α-Syn were measured and compared between phosphate-buffered saline (PBS)-treated neurons *vs.* neurons treated with WT or Ni PFFs. Data revealed a significant reduction of α-Syn protein in the soluble fraction of WT but not Ni PFFs-exposed neurons (Fig. 3E). This finding likely reflects a shift in endogenous α-Syn from the soluble to insoluble fraction during the seeding process, which was evident in WT PFFs-treated neurons and indicated aggregation. These results also confirmed the HCA analyses showing that significantly less seeding occurred after addition of Ni PFFs. Next, in the insoluble fractions of the PFFs-treated neurons, both monomeric and HMWs species (∼23, 37, 40, and 50 kDa) were detected on D10 and D21 by a pan-synuclein antibody (SYN-1) (Supplemental Fig. S7B). Strikingly, larger (>110 kDa) SDS-resistant aggregates were more effectively detected by the total α-Syn antibody (SYN-1) in the insoluble fractions of Ni as compared to WT PFFs-treated neurons (Supplemental Fig. S7B). The SYN-1 antibody detects both human PFFs and mouse endogenous α-Syn contained in seeded aggregates in the insoluble fraction. Therefore, this antibody does not allow for distinguishing whether these HMWs are newly formed fibrils or simply result from the cross-linking of nitrated PFFs that generate HMWs species. Thus, the mouse-specific total α-Syn (Glu105) antibody (Fig. 3F) and the pS129 (MJF-R13) antibody (Fig. 3G) were used to discriminate the newly formed α-Syn aggregates generated from the endogenous mouse α-Syn expressed in the primary neurons from the exogenously added human PFFs. As the latter undergo C-terminal cleavage, they do not undergo phosphorylation at S129^32, 46, 50^ and cannot be detected by the mouse-specific-Glu105 antibody. Consistent with ICC and HCA measurements, the results confirmed that newly formed fibrils were barely detected in the insoluble fraction of the nitrated PFF-treated neurons on D10. Despite a modest increase in seeding between D10 and D21 in the nitrated PFFs-treated neurons (Fig. 3F–G), levels of both total insoluble α-Syn (Glu105) and total pS129-positive (MJF–R13) α-Syn remained much lower in Ni PFFs-exposed than WT PFFs-treated neurons (Fig. 3F–G). Similar results were obtained with different batches of unmodified and nitrated PFFs prepared independently (batches #3; Supplemental Fig. S4 and S8). Together, these results demonstrated that post-fibrillization nitration significantly reduced the formation of insoluble aggregates containing endogenous α-Syn after the addition of α-Syn PFFs to primary neurons, and showed that this effect was due to an intrinsically reduced ability of nitrated fibrils to seed α-Syn aggregation.

### Nitration of α-Syn PFF induces their fragmentation and alters their stability in a site-specific manner

Fibril length has been established as one of the key determinants of α-Syn PFFs seeding activity^27, 28^. Therefore, we sought to investigate the effect of nitration on the length of PFFs. We first measured the length of unmodified and nitrated α-Syn fibrils by TEM. As sonication breaks down fibrils into shorter filaments, these studies were performed using non-sonicated fibrils. TEM images, taken immediately after nitration and after quick removal of the nitrating agent (30 min), clearly showed that nitrated fibrils were already significantly shorter in length than the unmodified fibrils (Fig. 4A). Indeed, the quantification of fibril length revealed that nitrated fibrils already displayed a shorter median (104 nm) and smaller range (24–525 nm) at 30 min (time point 0) after nitration (nitrated fibrils; Fig. 4A). Conversely, unmodified fibrils presented a much wider range of length (0.16–1628 nm), with a median length of 384 nm (unmodified fibrils; Fig. 4A). These findings suggest that nitration of unmodified α-Syn itself induces fibril fragmentation. Next, to assess if nitration of α-Syn induces further fragmentation or remodeling of the fibrils over time, we monitored changes in the fibril size of unmodified and nitrated fibrils over 35 days (37°C, 300 rpm) (Supplemental Fig. 2B). On average, the median length of the unmodified fibrils decreased by about 67% after 21 days compared to time point 0 (from about 380 to 126 nm; Fig. 4A), supporting previous findings of the continuous shortening of fibrils over time^51, 52^. Conversely, the length of the nitrated fibrils at time 0 (median 104 nm) changed more moderately after 35 days (to a median of about 69 nm), dropping by about 34% in 35 days. Furthermore, the length of both unmodified and nitrated fibrils declined following distinct patterns. While the median length of unmodified fibrils dropped significantly by 41% to 226 nm within the first 7 days, the length of the nitrated fibrils remained essentially unchanged for 14 days. Between 7 and 14 days, the length of the unmodified fibrils also did not change significantly. Between 14 and 35 days, both fibril types underwent significant further shortening, by 45% for the unmodified and 34% for the nitrated fibrils. Taken together, these results suggested that nitration induced immediate fragmentation of α-Syn fibrils, whereas unmodified fibrils underwent slower fragmentation or fibril shortening over time, possibly due to continuous disassociation processes.

**Figure 4:**
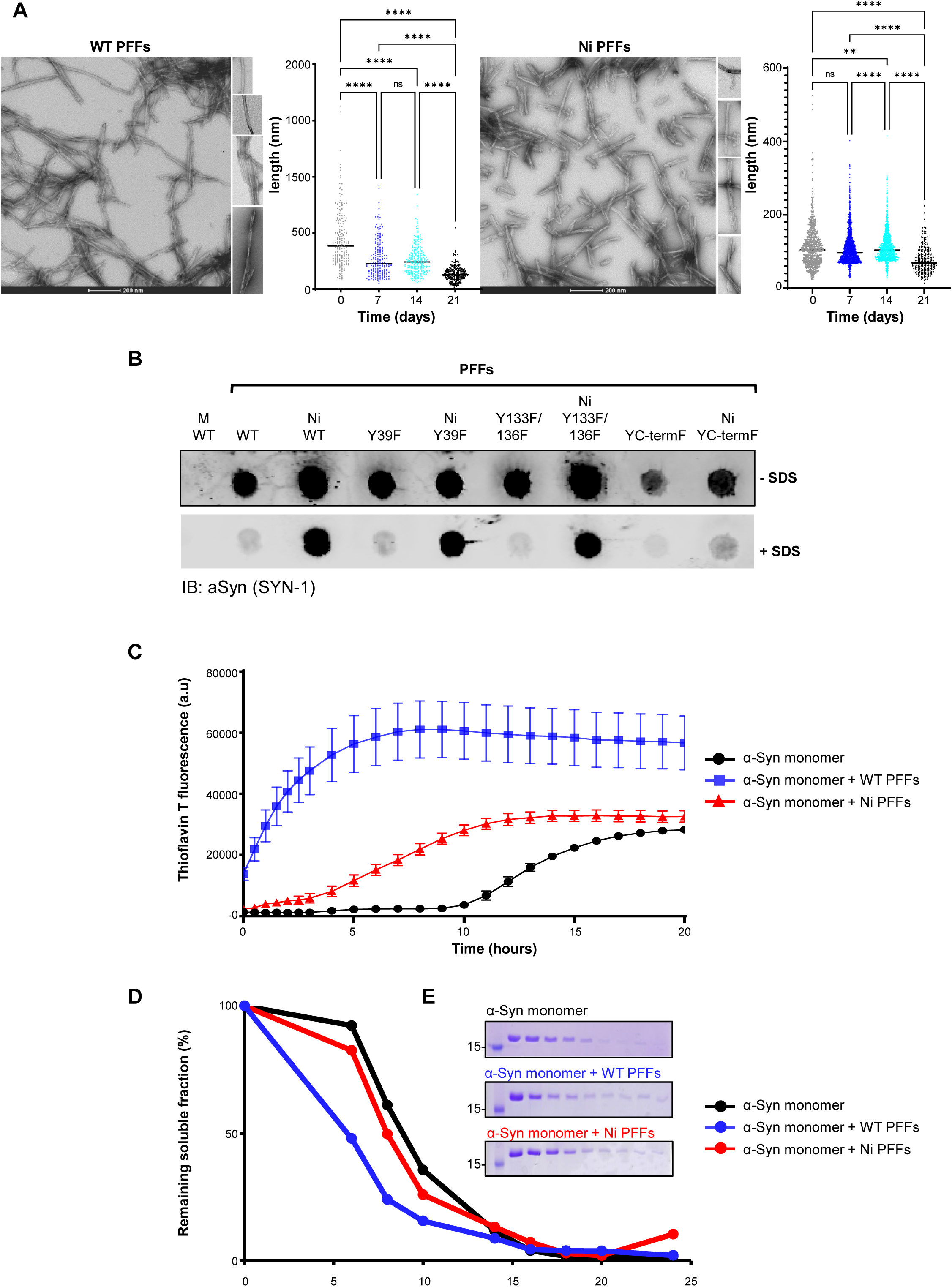
Nitrated fibrils were shorter in length compared to unmodified fibrils, and showed reduced seeding activity. **A.** TEM images of unsonicated fibrils before (left image) or after (right image) nitration, with the corresponding length of the fibrils on their side. **B.** Filter trap assay for monomeric, fibrillar WT and Y->F mutants α-Syn, and their respective nitrated forms in the presence or absence of denaturing buffer (SDS). **C.** Fluorescent ThT aggregation assay and **D**. Sedimentation assay of monomeric α-Syn (black dots/line), monomeric α-Syn + unmodified fibrils (blu dots/line), and monomeric α-Syn + nitrated fibrils (red dots/line). **E.** Representative SDS-PAGE of the remaining soluble protein at different incubation time points. Monomeric α-Syn (upper blot), monomeric α-Syn + unmodified fibrils (central blot), and monomeric α-Syn + nitrated fibrils (lower blot). Scale bars = 200 nm. The images are qualitative.

Nitration and intra-molecular dityrosine formation in the flexible C-terminal domain is expected to stabilize the monomers within the fibrils and prevent or slow their disassociation. Furthermore, given that residue Y39 is closer to the core of α-Syn fibrils, one would expect that nitration at Y39 might result in more fragmentation of α-Syn PFFs, whereas blocking nitration at Y39 might lead to the formation of larger fibrillar structures or fibrillar networks that are stabilized by dityrosine cross-linking. To test this hypothesis, we used the filter trap assay to compare the retention of unmodified and nitrated α-Syn PFFs derived from WT or N-and C-terminal tyrosine mutants [YC-termF(Y125F/Y133F/Y136F), Y133F/Y136F, or Y39F] in the absence and presence of SDS, using the SYN-1 total α-Syn monoclonal antibody (Fig. 4B). As expected, when WT and Y->F mutated α-Syn PFFs were diluted in a non-denaturing buffer, only the monomeric recombinant protein passed through the membrane, whereas all PFFs, independent of nitration status, were retained in the membrane. However, when the samples were diluted in SDS-containing denaturation buffer, only PFFs that could be nitrated at the C-terminal of α-Syn protein (NitroWT; NitroY39F; Nitro Y133F/Y136F) were retained in the membrane, indicating intra-and inter-fibril cross-linking-mediated stabilization of the fibrils. By contrast, WT PFFs or YC-termF(Y125F/Y133F/Y136F) PFFs that could only be nitrated at Y39 were not retained on the membrane, consistent with Y39 nitration-mediated destabilization/fragmentation of the fibrils. These results suggest that the nitration-induced fragmentation of the fibrils appears to be driven by the nitration of Y39.

### Post-fibrillization nitration of α-Syn PFFs reduces their seeding activity *in vitro* in a PFF length-independent manner

Several *in vivo* studies have shown that shorter fibrils are more effective in seeding α-Syn fibrillization in neurons and *in vivo* due to the increased number of extension sites per weight of fibril material, compared with long fibrils^53, 54^. Our observation that post-fibrillization nitration modifies fibril length thus prompted us to further investigate the mechanism by which the nitration of α-Syn fibrils may alter their seeding activity, especially since we observed minimal seeding activity by nitrated α-Syn PFFs *in vivo* and in neuronal models of LB formation. Therefore, we compared the seeding capacity of nitrated WT and YF mutated forms of α-Syn fibrils versus unmodified fibrils *in vitro* using the Thioflavin T (ThT) assay (Fig. 4C and Supplemental Fig. S9). Figure 4C shows that both unmodified and nitrated fibrils efficiently seeded the aggregation of unmodified monomeric α-Syn *in vitro*, although the nitrated fibrils exhibited slightly reduced seeding activity. The kinetics of aggregation of WT monomeric α-Syn were in accordance with previous studies^3, 26, 28, 30^, with α-Syn fibrillization reaching a plateau after >24 h (Fig. 4C, black line). When WT monomeric α-Syn was co-incubated with WT fibrils as seeds, the kinetics of aggregation, as expected, was faster (i.e., the lag time was shorter) (Fig. 4C, blue line). Under the same conditions, the slope of ThT fluorescence was initially less steep for nitrated fibrils (Fig. 4C, red line). This finding indicates that the aggregation kinetics are slower for nitrated fibrils shortly after the start of the seeding experiment. However, ThT fluorescence also steeply increased for the nitrated fibrils with about a 1 h delay compared to the unmodified fibrils. As a result, both nitrated and unmodified fibrils reached plateau phases of ThT fluorescence after about the same time (∼10 h). Similar results were obtained when the seeded aggregation reactions were assessed by quantifying the amount of remaining soluble α-Syn using the well-established sedimentation assay (Fig. 4D, E^33^). The reduced seeding activity of the nitrated fibrils, despite the fact that they were shorter compared to fibrils formed by WT protein, could be explained by strong intramolecular cross-linking, which renders the nitrated α-Syn PFFs less dynamic with reduced monomer release. These results suggest that the dynamic properties of the fibrils, rather than simply their length, is an important determinant of fibril seeding capacity *in vitro*. Alternatively, it is possible that extensive nitration of the C-terminal domain, which coats the surfaces of the fibrils, may alter their ability to mediate seeding via secondary nucleation events.

Based on these assumptions, we hypothesized that fibrils, which are more prone to fragmentation (e.g., α-Syn fibrils nitrated at Y39; Fig. 4) and heavily intramolecularly cross-linked fibrils (e.g., fibrils nitrated at C-terminal tyrosines) would display different seeding capacities. To test this hypothesis, we measured the seeding capacity of α-Syn fibrils specifically nitrated at all C-terminal tyrosine residues (Y39F fibrils; Supplemental Fig. 9A) or that are partially nitrated at Y39 (YctermF fibrils; Supplemental Fig. 9B) compared to the unmodified WT fibrils. α-Syn fibrils nitrated specifically at Y39 displayed a similar seeding capacity as the unmodified fibrils (Supplemental Fig. 9B). This could be explained by the fact that Y39, within PFF, is only partially nitrated. By contrast, nitration of C-terminal tyrosine residues showed a tendency towards decreased seeding (Supplemental Fig. 9A).

## Discussion

Several lines of evidence point to the complex interplay between oxidative and nitrative stress and protein aggregation and neurodegeneration in several NDDs. However, the relative contribution and differential role of oxidative stress at different stages of the development and progression of NDDs remains poorly understood^55^. One major hallmark of oxidative stress in NDDs is the preferential accumulation of nitrated misfolded proteins in the proteinaceous pathological inclusions found in the brains of affected patients^2, 7, 56^. This has led to the suggestion that oxidative modifications of aggregation-prone proteins may play causative roles in the initiation and/or progression of protein misfolding, aggregation, pathology spreading, or neurodegeneration in NDDs^20, 57^. Therefore, a large body of literature on the role of protein nitration in NDDs is dominated by studies exploring the pathogenic role of nitration and its role in modifying the behavior and function of aggregation-prone proteins linked to NDDs at the level of the native monomeric form of these proteins.

In this study, we explored for the first time the role of post-aggregation nitration in regulating key processes that drive pathology formation and spreading in PD, namely the ability of α-Syn fibrils to induce the aggregation of monomeric α-Syn *in vitro*, LB-like inclusion formation in neuronal cultures, and pathology formation and spreading in the mouse brain. This study was motivated by several lines of evidence, including work from our laboratory showing that: 1) non-specific nitration of α-Syn monomers stabilizes non-fibrillar aggregates and inhibits the formation of α-Syn fibrils^26–30^ (Fig. 5), which are the dominant aggregated form of α-Syn found in LBs and other types of Lewy pathologies in PD and other synucleinopathies; 2) site-specific nitration of α-Syn at Y39 or Y125 did not alter its aggregation kinetics, but did change the morphological properties of α-Syn in fibrillar aggregates^30^; and 3) several α-Syn PTMs that are enriched in Lewy pathologies are not required for inducing α-Syn aggregation and inclusion formation. Based on these premises, we hypothesized that the nitration of tyrosine residues might not be the primary driver of α-Syn fibrillization *in vivo* but could still modulate the pathogenicity of α-Syn by influencing which fibril strains are formed or by modulating the seeding activity and spreading properties of the fibrils. We postulated that post-aggregation nitration could either be pathogenic (i.e., enhance α-Syn seeding activity and spreading or toxicity) or protective (i.e., suppresses the pathogenic activity of α-Syn PFFs); in particular, a detoxifying effect may arise from rendering α-Syn fibrils seeding incompetent. An attractive feature of such a protective mechanism is that its beneficial effects may be independent of the nature of specific fibril strains since three of the four tyrosine residues in α-Syn are located at the extreme C-terminus of the protein. Indeed, all cryo-EM structures of α-Syn fibrils, with only one exception^58^, have shown that this protein region remains flexible and accessible for chemical and enzymatic-mediated modifications^35, 36, 57, 59^. Consistent with this hypothesis, we showed that nitration of both mouse and human α-Syn fibrils nearly abolishes their seeding activity, despite large differences in the morphological properties of the two PFF preparations (Supplemental Fig. S4). Studies are underway to determine if nitration of fibrils represents a common mechanism for inducing the fragmentation and reducing the sedding activity of other aSyn fibrils strains and fibrils derived from other amyloid proteins linked to other NDDs (e.g. Tau).

**Figure 5:**
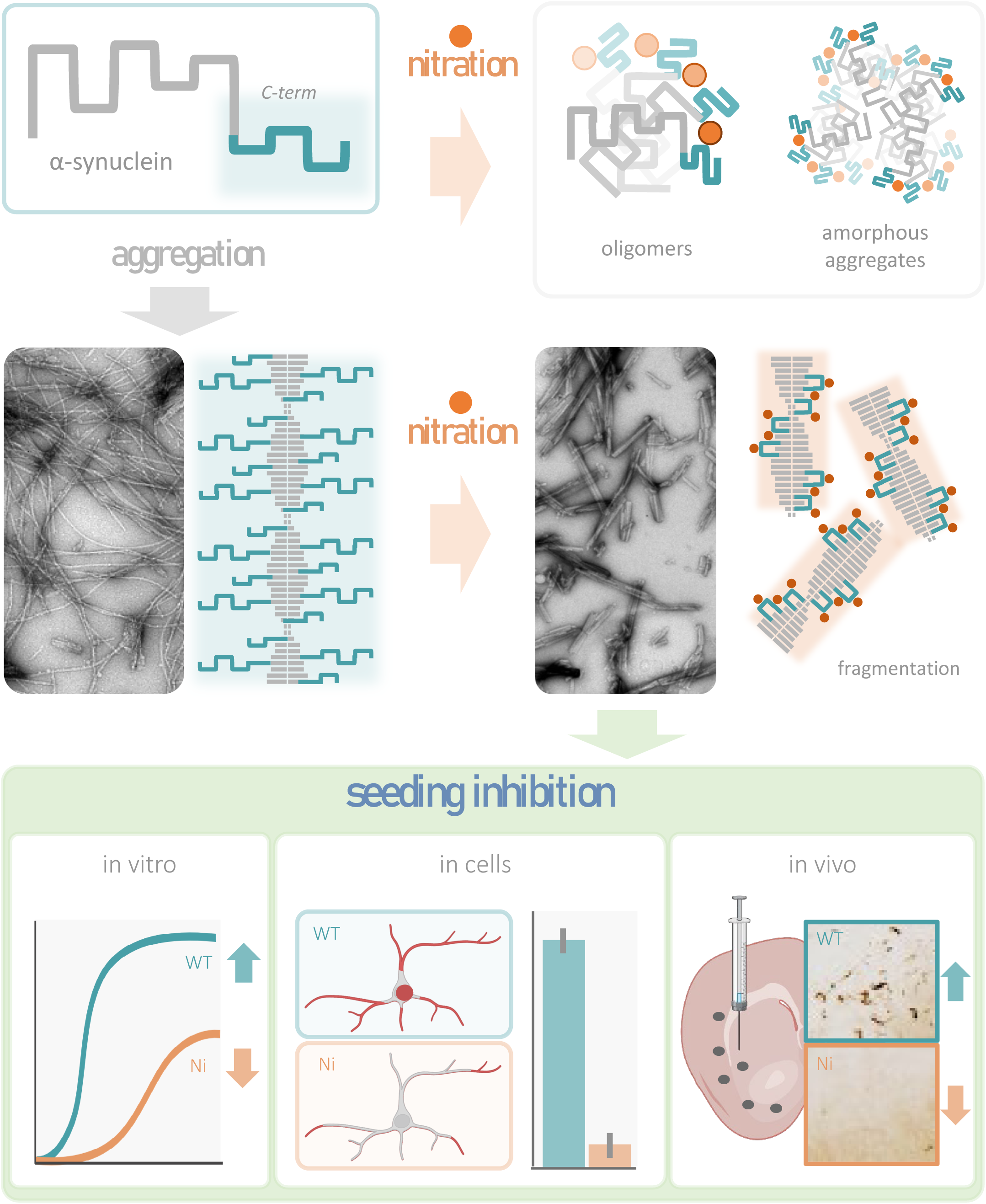
Schematic depictions illustrating our current understanding of the impact of nitation of aSyn monomers and summarizing our results on the effect of post-fibrillization nitration on α-Syn PFFs. Nitration of aSyn monomers results in the formation of non-fibrillar and amorphous aggregates that do not convert to fibrils. Nitration of α-Syn PFFs induces their fragmentation, modifies their surface properties, and inhibits their seeding in vitro, in primary neurons and in vivo.

Our comparative biophysical and seeding studies on unmodified and nitrated α-Syn fibrils provide some hints into possible mechanisms by which the nitration of PFFs could modulate their propagation and pathogenicity. First, our MS analyses revealed that the chemical nitration of fibrillar α-Syn resulted in the complete modification of all three C-terminal tyrosine residues and partial nitration of Y39 (Fig. 1A). This is consistent with the Y39 residue being located in the core of the helix facing the outside of the fibrils and being, therefore, only partially accessible for modifications (Fig. 1 and Supplemental Fig. 3). By contrast, the C-terminal region of α-Syn fibrils is not resolved in most cryo-EM structures of α-Syn fibrils, suggesting that this region is more disordered, flexible, and thus possibly more accessible for modifications (e.g., by nitrogen dioxide groups). Second, nitration-induced cross-linking of α-Syn results in more stable and less dynamic nitrated α-Syn PFFs than unmodified PFFs. It is well established that nitration of α-Syn induces the formation of dityrosine cross-linked α-Syn oligomers *in vitro*^30^. Several studies have also reported the detection of dityrosine cross-linked α-Syn in the brains of PD patients^13, 30^. In the case of intrinsically disordered proteins like α-Syn, these cross-links may strongly influence the ensemble of conformations adopted by the polypeptide chain^60^. The occurrence of dityrosine formations on PFFs could lead to their stabilization by promoting the cross-linking of monomers from different filaments. Alternatively, it is possible that extensive nitration of the C-terminal domain, which coats the surfaces of the fibrils, may alter their ability to mediate seeding via secondary nucleation events and, by doing so, may affect fibril-mediated pathology formation and propagation (Fig. 5).

In this study, our extensive analyses of α-Syn fibrils before and after nitration using TEM and ssNMR revealed that post-fibrillization nitration of α-Syn *in vitro* did not substantially change the morphology, diameter, or structure of α-Syn fibrils. The nitrated fibrils showed a predominantly rod-like structure and were indistinguishable from control fibrils. By contrast, we observed that nitration of the PFFs led to their immediate fragmentation, as evidenced by a significant shortening of the length of these fibrils, which persisted over several weeks. Taking advantage of the site-specific replacement of tyrosines with phenylalanine that enabled site-specific nitration, we identified nitration at Y39, which is located in the core of most fibril structures, as the key determinant of PFF fragmentation *in vitro* (Supplemental Fig. 9). Initially, these observations led us to speculate that fibril fragmentation could increase the number of seeding competent fibrils, which could lead to enhanced α-Syn PFF seeding activity and potentially pathology spreading. To test this hypothesis, we investigated the seeding activity of nitrated α-Syn fibrils *in vitro*, in neuronal cultures, and in a mouse model of seeding-mediated α-Syn pathology formation. The results from these studies demonstrated that post-fibrillization nitration of WT α-Syn PFFs increased their stability, significantly reduced their seeding capacity *in vitro*, and nearly abolished their seeding activity in primary neurons and mice.

The similar *in vitro* seeding properties of fibrils nitrated at only Y39 residues (Y125F/133F/136F-α-Syn PFFS) and unmodified fibrils (Supplemental Fig. 9), despite differences in fibril lengths between the two PFFs preparations, suggests that changes in the stability, dynamic properties, and biochemical surface properties of α-Syn PFFs, linked to the nitration of C-terminal tyrosine residues, is the primary contributor to the seeding inhibitory effects of the nitrated PFFs. Our observation that the effects of post-fibrillization nitration on α-Syn PFF seeding activity were more pronounced in primary neuron cultures and *in vivo*, compared to cell-free systems, suggests that other cellular factors or molecular interactions play critical roles in regulating α-Syn seeding and fibril growth in neurons. It is also plausible that nitration disrupts the interaction of fibrils with other proteins, membranes, or organelles that enhance the seeding activity of the fibrils^61^. Future studies are needed to explore the impact of post-fibrillization PTMs on the interactome of α-Syn oligomers and fibrils.

One possible explanation for the lack of seeding activity of nitrated PFFs in neurons or *in vivo* is the reduced uptake or differentiation processing of fibril post-internalization into neurons. However, our results demonstrate that this is not the case. The nitration of α-Syn PFFs did not influence their internalization or processing in neurons, namely C-terminal cleavage. The latter also suggests that the nitrated C-terminal domain of PFFs remains dynamic and accessible to proteases.

Several α-Syn PTMs, in addition to nitration^2, 18, 19, 62^, are consistently associated with LB and pathological inclusions^63–67^. Our current work and previous studies from multiple groups demonstrate that PTMs serve as key regulators of α-Syn oligomerization, seeding and fibrillization, LB formation, and pathology propagation. It is likely that different types of PTMs have differential effects on the α-Syn fibril interactome, seeding activity, and clearance and that targeting post-aggregation PTMs may provide new opportunities for modulating α-Syn pathology formation and neurotoxicity. Dissecting the role of PTMs in these processes will require more insights into 1) the role of PTMs in regulating the interactome of α-Syn fibrils; 2) the key enzymes and cellular processes that regulate α-Syn PTMs in health and disease; 3) and the development of novel tools to induce or regulate α-Syn PTMs at different stages of LB formation with spatial and temporal resolution.

### Pathogenetic and therapeutic implications

To the best of our knowledge, this is the first study showing that post-fibrillar modifications of α-Syn may provide a therapeutic approach for potentially slowing down disease progression. This modification may include “targeted” nitration or other PTMs that exert similar effects on the biochemical and surface properties of α-Syn fibrils.

To date, the majority of therapeutic approaches targeting oxidative stress in PD have focused on attenuating oxidative stress and/or inhibiting α-Syn nitration. Indeed, a significant body of experimental evidence supports a deleterious role of oxidative stress in promoting α-Syn pathology and, in particular, its aggregation and inter-neuronal spreading^21, 22, 68^. The work presented here significantly advances our understanding of the relationship between oxidative stress and pathogenetic processes involving α-Syn. By demonstrating that nitration of α-Syn fibrils markedly attenuates further protein aggregation and seeding-mediated pathology spreading, our current findings support an intriguing new concept and indicate that oxidative stress may play divergent roles at different stages of α-Syn pathology and disease development. In particular, oxidative and nitrative α-Syn modifications may contribute to toxic changes at early pathogenetic stages when a deleterious role of oxidative stress could be primarily mediated *via* modifications of relatively soluble α-Syn aggregates, such as oligomers^21, 22, 68^. A different scenario would arise after the formation and accumulation of mature α-Syn fibrils and α-Syn-containing inclusions. Under these conditions, post-aggregation nitration is likely to exert protective effects, inhibiting further progression of α-Syn pathology. The notion of different roles of oxidative/nitrative reaction in PD pathogenetic processes raises at least two important considerations. The first consideration relates to the consequences of Lewy body accumulation in PD and the hypothesis that this accumulation may represent a protective mechanism that segregates toxic α-Syn species and prevents their toxic interactions with other cytosolic molecules and cellular organelles^46, 68^. Our present findings are consistent with this hypothesis and, more importantly, reveal a specific mechanism by which the formation of Lewy inclusions may “buffer” the burden of oxidative α-Syn species and counteract pro-aggregation and seeding reactions. The second corollary of our findings revealing a protective role of oxidative reactions concerns the possibility that mimicking the biochemical effects of post-aggregation nitration may provide a new strategy for therapeutic intervention in PD and other synucleinopathies. Our proof-of-concept evidence points to mature α-Syn fibrils as important therapeutic targets and indicates that PTMs of these fibrils could effectively lower their pathogenicity and counteract α-Syn-dependent disease progression.

It has been proposed that α-Syn and other amyloid proteins form fibrils of different strains, and the possibility that differences in α-Syn fibril strains could contribute to the clinical heterogeneity of PD and other diseases has been noted. Therefore, it is plausible that variation in the biochemical and structural properties of α-Syn aggregates in different brain regions and between different affected individuals could explain the failure of current therapies, such as immunotherapies, that have not accounted for such pathological heterogeneity. One possible approach may involve the development of therapies targeting different strains. Unfortunately, this is not currently possible due to the lack of conditions that allow reproducing the structures and biochemical properties of α-Syn fibrils found in the brains of patients with PD, MSA, or LBD. Our work points to a new approach that could pave the way for novel therapies that address α-Syn pathological diversity because the effect of post-aggregation nitration of α-Syn should be insensitive to differences in fibril morphologies and strains. This is because the region harboring three key nitratable α-Syn residues decorates the surfaces of all α-Syn fibril polymorphs, thus remaining exposed or accessible to modifications (Fig.5). In fact, our study shows that post-fibrillization nitration nearly abolished the seeding activity of human as well as mouse α-Syn PFFs (Supplemental Fig. 10), even though these two preparations exhibited distinct morphological and seeding activities.

Together, our findings suggest that the targeted nitration of fibrils may represent an effective strategy to prevent the accumulation and spreading of pathology involving α-Syn fibrils of different morphologies/strains. We recognize that elevating oxidative stress is unlikely to emerge as a viable therapeutic strategy. However, identifying mechanisms/pathways that could be activated to selectively induce α-Syn nitration or other PTMs that exert similar effects on pathogenic α-Syn fibrils could open the door to new PTM-dependent disease-modifying strategies for PD and other synucleinopathies. The diminished pathogenic properties of nitrated α-Syn fibrils show that not all fibrils found in the brain are pathogenic and suggest that simple quantification of α-Syn pathology levels in the brain may not be the best way to investigate the relationship between pathology formation and disease development and progression. The presence of a mixture of different levels of pathogenic and non-pathogenic α-Syn could partially explain the often-reported lack of correlation between α-Syn pathology burden and disease progression and severity. We postulate that the pattern of PTMs on pathological aggregates may be a crucial factor in determining their toxicity and impact on neurodegeneration. Future studies using antibodies against multiple α-Syn PTMs, rather than relying on only pS129 antibodies, may provide a more reliable approach to dissecting the relationship between α-Syn pathology formation and neurodegeneration in NDDs and preclinical models of synucleinopathies. Finally, we believe that our findings are likely to bear pathogenic and therapeutic implications relevant for other proteinopathies that, similar to PD, are characterized by the accumulation of pathological aggregates with extensive PTMs. Therefore, further investigations are warranted to elucidate the physiopathological role of nitration and other PTMs in diseases such as Alzheimer’s disease and transactive response DNA-binding protein 43 proteinopathies.

## Material and Methods

### Antibodies

All of the primary and secondary antibodies used in this study are described in Figure S1.

### Recombinant overexpression and purification of human WT and mutated α-Syn

Recombinant overexpression and purification of human WT or Y->F mutants (Y39F; Y133F/Y136F; and Y125F/Y133F/Y136F) α-Syn proteins were performed as previously described^33^. pT7-7 plasmids encoding human WT or mutated α-Syn were used for transformation in BL21 (DE3) *Escherichia coli* cells on ampicillin agar plates. For each protein, a small-scale culture was prepared by transferring a single colony to 200 mL Luria broth (LB) medium containing 100 μg/mL ampicillin (AppliChem) and incubated overnight at 180 rpm at 37°C. The next day, a large-scale culture was prepared by inoculating the pre-culture into 6 L LB medium containing 100 μg/mL ampicillin. When the absorbance at 600 nm reached 0.4–0.6, α-Syn protein expression was induced by the addition of 1 mM 1-thio-β-d-galactopyranoside (AppliChem), and the cells were further incubated at 180 rpm at 37°C for 4–5 h. After this incubation time, the absorbance at 600 nm typically reached 1–1.2, after which the cells were harvested by centrifugation at 4,000 rpm for 30 min at 5°C using the JLA 8.1000 rotor (Beckman Coulter). The harvested pellets were stored at −20°C until use. Cell lysis was performed by dissolving the bacterial pellet in 100 mL of 40 mM Tris-hydrochloride (HCl; Sigma-Aldrich) buffer (pH 7.5) containing protease inhibitors, 1 mM ethylenediaminetetraacetic acid (Sigma-Aldrich), and 1 mM phenylmethane sulfonyl fluoride (PMSF; Applichem), followed by ultrasonication (Sonic Vibra Cell, Blanc-Labo) in the following conditions: 8 min; cycle: 30 s on, 30 s off; amplitude 70%. After lysis, centrifugation at 12,000 rpm at 4°C for 30 min was performed to collect the supernatant in 50 mL Falcon tubes, which were placed in boiling water (100°C) for ∼15 min to allow the heat-sensitive proteins to precipitate. This solution was subjected to another round of centrifugation at 12,000 rpm at 4°C for 30 min, and the obtained supernatant was filtered through 0.45-μm filters and injected into a sample loop connected to HiPrep Q Fast Flow 16/10 column (Sigma-Aldrich). The protein was eluted using 40 mM Tris-HCl, 1 M sodium chloride (NaCl) buffer (pH 7.5) from a gradient of 0% to 70% at 3 mL/min. All fractions were analyzed by SDS-PAGE, and the fractions containing pure α-Syn were loaded on a reverse-phase high-performance liquid chromatography (HPLC) C4 column (PROTO 300 C4 10 μm, buffer A, 0.1% trifluoroacetic acid [TFA] in water; buffer B, 0.1% TFA in acetonitrile; Higgins Analytical). The protein was eluted using a gradient from 35% to 45% buffer B over 40 min (15 mL/min). The purity of the elution from HPLC was analyzed by ultra-performance LC (UPLC) and ESI-MS. Fractions containing highly purified α-Syn were pooled, snap-frozen, and lyophilized.

For some experiments, human WT or mutated α-Syn proteins purified without TFA were used. In those cases, instead of being loaded on a reverse-phase HPLC C4 column, the fractions containing purified α-Syn were pooled and centrifuged in 30 kDa MW cut-off (MWCO) tubes at 4,000 g for 15 min at 4°C to concentrate the protein and remove any impurities less than 30 kDa. Then the retentate was purified by size exclusion chromatography (HiLoad 26/600 Superdex 200 pg), and the fractions of interest were analyzed by SDS-PAGE. The fractions containing pure α-Syn were dialyzed against distilled water using dialysis tubes (12–14 kDa MWCO) overnight at 180 rpm at 4°C to remove excess salts. Then the proteins were analyzed by UPLC and ESI-MS, and only the pure α-Syn fractions were pooled, snap-frozen, and lyophilized.

### Preparation of human WT and mutated Y->F α-Syn fibrils for *in vitro* and *in vivo* studies

WT and Y->F mutants (Y39F, Y133F/136F, and Y125F/133F/136F) α-Syn fibrils were prepared by dissolving 4 mg lyophilized recombinant α-Syn in 600 μL PBS, and the pH was adjusted to ∼7.2–7.4 using a 1 M sodium hydroxide solution. The solution was transferred to Spin-X centrifuge tube filters (Costar 0.22 μm polypropylene) and centrifuged at 5,000 *g* for 3 min at 4°C. After centrifugation, a small aliquot of the protein was subjected to ESI-MS to verify the MW and identify possible impurities. Another aliquot was mixed with 4X Laemmli buffer (10% SDS, 50% glycerol, 0.05% bromophenol blue, 1M Tris-HCl pH 6.8 and 20% β-mercaptoethanol) and stored at −20°C for subsequent SDS-PAGE analyses representing the starting monomeric protein. The concentration of monomeric protein was assessed by measuring the absorbance at 280 nm on a Nanodrop 2000 spectrophotometer (Thermo Fisher Scientific) and using an extinction coefficient of 5,960 M^−1^ cm^−1^ predicted using the sequence (ProtParam, ExPASy). The remaining protein was transferred to black screw cap tubes and incubated in a shaking incubator at 37°C and 1,000 rpm for 5 days, after which the solution was centrifuged at 100,000 *g* for 30 min at 4°C to separate the monomers and oligomers that did not undergo fibrillation from the insoluble fibrils^33^. Fibrils were resuspended in 600 μL PBS by vigorous pipetting and vortexing. The formation of fibrils was assessed by TEM. The different species (monomers/oligomers/fibrils) in the solution were quantified by Coomassie staining as previously described^33^.

For the preparation of sonicated seeds for neuronal and *in vivo* seeding studies, WT and Y->F mutant α-Syn fibrils were subjected to sonication for 20 s, at 20% amplitude 1 s pulse on and 1 s pulse off (Sonic Vibra Cell; Blanc-Labo). The amount of monomer and oligomer release from sonicated fibrils was quantified by filtration as previously reported^33^. In addition, the structure of the fibrils was monitored by TEM, and the amount of released monomers and oligomers after sonication was quantified by SDS-PAGE and Coomassie staining. Only seeds with an average length between 50 and 100 nm were used for subsequent experiments.

### TEM

To examine the ultrastructure of α-Syn fibrils, negative staining electron microscope images were taken using Formvar/carbon-coated 200-mesh copper grids (Electron Microscopy Sciences). Activated grids were loaded with 5 μL sample for 30 s, washed three times with ultrapure water, and then negatively stained with 1% uranyl acetate for 1 min. Excess liquid was removed, and grids were allowed to air dry. Imaging was conducted on the Tecnai Spirit BioTWIN electron microscope at 80 kV acceleration voltage and equipped with a digital camera (FEI Eagle; FEI). A total of 5–10 images for each sample were chosen, and the length and width of the fibrils were quantified using ImageJ software (RRID:SCR_001935; National Institutes of Health). For each sample ∼800–1000 fibrils were quantified.

### Post-fibrillization nitration of WT and Y->F mutated α-Syn

Post-fibrillization nitration of WT or Y->F mutants α-Syn was induced according to a previously published protocol^5^. Briefly, α-Syn fibrils were incubated with 10 equivalent (eq.) of tetranitromethane (TNM, 10% in ethanol) per tyrosine residue to protein (2 mg/mL) in filtered Tris-buffered saline (TBS) (for *in vitro* experiments) or in sterile 1X PBS buffer, pH 7.4 (Gibco) (for cellular and *in vivo* experiments). The reaction mixture was stirred vigorously for 30 min at 37°C, after which an additional 10 eq. of TNM per tyrosine residue was added, and the reaction was stirred under the same conditions for 90 min. Following nitration, the fibrils were washed three times with TBS to remove the excess TNM by using the 100 kDa MWCO filters (Millipore). The centrifugation steps were performed at 20,000 g for 30 min at 4°C. The nitrated fibrils were collected on the membrane during the first two washing steps. For the last washing step, 50 µL TBS was added to the Eppendorf tube, the filter containing the nitrated fibrils was carefully turned upside-down, and 100 µL TBS was slowly added. After centrifugation (5,000 g, 5 min, 4°C), the pelleted nitrated fibrils were resuspended by vigorous pipetting and vortexing. Then the nitrated fibrils were boiled for 10 min at 95°C and spun down, after which the supernatant was subjected to ESI-LC/MS.

### ESI-LC/MS analysis

MS analyses of proteins were performed on the Agilent 1100 Series quadrupole mass spectrometer equipped with an electrospray ion source, and the spectrometer was operated in positive ion mode as previously described^3^. Before analysis, fibrillary proteins roughly at a concentration of 0.5–1 μg were boiled for 10 min at 95°C and centrifuged for 30 min at 13,000 rpm. The monomeric form of the protein was injected into the column and eluted from 5% to 95% of solvent B against solvent A with a linear gradient. The solvent composition of solvent A was 0.1% formic acid in ultra-pure water, and that of solvent B was 0.1% formic acid in acetonitrile. MagTran software (Amgen Inc.) was used for charge state deconvolution and MS analysis.

### Remodeling of WT and Y->F mutants α-Syn

Unmodified or Y->F mutants α-Syn were prepared as above and not subjected to sonication. Briefly, each type of fibril was divided into two aliquots. One aliquot was subjected to post-fibrillization nitration as described above, and the second aliquot was kept on ice. After the removal of excess TNM from the nitrated fibrils, all of the fibrils were diluted at a final concentration of 20 μM and incubated at 37°C continuously. The fibrils were gently shaken for 15 min per hour at 300 rpm to avoid fibril accumulation on the bottom of the tube. To follow the structure of the fibrils over time, aliquots from all the samples were taken at time 0, and after 7, 21, 28, and 35 days. From all aliquots, TEM grids were prepared immediately, and then 5–10 images for each sample were taken and analyzed. The length and the width of the fibrils were measured as described above.

### Proteinase K resistance of WT and Y->F mutants α-Syn

Unmodified or nitrated fibrils used for proteinase K (ProK) digestion were prepared as above and not subjected to sonication. Briefly, all fibrils at a concentration of 1 mg/mL were incubated with various concentrations (0, 0.5, 1.0, 1.5, and 2.0 μg/mL) of ProK in 10 mM PBS (pH 7.4) at 37°C for 30 min. Following incubation, the reaction was quenched by the addition of 5 μL of 5X Laemmli buffer to the reaction mixture. The samples were boiled for 8–10 min at 95°C and loaded onto precast 12% Bis-Tris gels (Bio-Rad) with MES SDS running buffer (Bio-Rad) and run at 120 V for 60 min. The SDS bands were visualized using Coomassie blue staining or WB. α-Syn bands were detected with SYN-1 monoclonal mouse antibody.

### Filter trap assay of WT and Y->F mutants α-Syn

Unmodified or nitrated fibrils used for the filter trap assay were prepared as above and subjected to sonication. Briefly, all fibrils were diluted to a final concentration of 10 µM in a buffer containing 1% SDS. The samples were loaded by vacuum filtration onto a 96-well dot blot apparatus (Bio-Rad) containing a cellulose acetate membrane (0.2 microns). Monomeric α-Syn was used as a control. The samples on the membrane were rinsed twice with 200 µL of 0.1% SDS buffer. After releasing from the blotting apparatus, the membrane was blocked for 1 h at room temperature (RT) with the Intercept (PBS) Blocking Buffer (LiCor), and immunoblot analysis was performed as described in the section below.

### ThT fluorescence-based aggregation kinetics *in vitro*

For ThT aggregation kinetics *in vitro*, unmodified or Y->F mutants α-Syn fibrils or their nitrated forms were used as seeds and subjected to sonication. Briefly, the fibrils were fragmented by sonication for 20 s at 20% amplitude, 1 s pulse on, 1 s pulse off (Sonic Vibra Cell; Blanc-Labo). The amounts of monomers and oligomers released from sonicated fibrils were quantified by a sedimentation-filtration assay as previously described^33^. Only seed preparations with <10% monomer release were used in our experiments. A master mix solution of 350 μL was prepared and contained WT α-Syn monomer at a final concentration of 20 µM, to which a final ThT concentration of 20 μM and 10% of pre-formed seeds were added. Each master mix solution was added to three individual wells in a black 96-well optimal bottom plate (Costar). The plate was sealed with Corning microplate tape and transferred to a Fluostar Optima microplate reader (BMG LABTECH), where continuous orbital shaking at 900 rpm and a temperature of 37°C were maintained for the duration of the experiment. ThT fluorescence was monitored every 300 s at excitation and emission wavelengths of 450 ± 10 and 480 ± 10 nm, respectively. The assay was stopped after 24 h, and the samples at the endpoint were analyzed by TEM, SDS-PAGE, and Coomassie staining and compared to the samples at the beginning of the ThT aggregation kinetics.

### Solubility assay

Similar to the ThT aggregation kinetics experiments, a master mix solution was performed containing a final concentration of 20 μM monomeric WT α-Syn to which 10% of unmodified or nitrated pre-formed seeds were added. Aliquots of the master mix solution were loaded onto a black 96-well optimal bottom plate (Costar), and the plate was sealed with Corning microplate tape and transferred to a Fluostar Optima microplate reader (BMG LABTECH), where similar parameters as the ThT aggregation kinetics experiments were applied. At different time points (0, 6, 8, 10, 14, 16, 18, 20, and 24 h), the plate reader was paused, and aliquots were taken from each well of the plate for TEM and SDS-PAGE analyses. For SDS-PAGE analysis, each sample was centrifuged at 100,000 *g* for 30 min at 4°C and analyzed after carefully removing the supernatant.

### Recombinant overexpression and purification of human ^13^C-^15^N labeled WT α-Syn for NMR studies

Recombinant overexpression and purification of human ^13^C-^15^N labeled WT α-Syn protein were performed as described above using cells that were grown in a minimal medium containing, as final concentrations, 1X M9 salts, 0.4% w/v [^13^C]-glucose, 2 mM magnesium sulfate, 0.1 mM calcium chloride, 0.03 mg/mL thiamine hydrochloride, and 50 μg/mL ampicillin. The 1X M9 salts were composed of sodium phosphate, monopotassium phosphate, NaCl, and [^15^N]-NH_4_Cl, pH 7.4 and autoclaved. The protein was purified using the HiPrep Q Fast Flow 16/10 column (Sigma-Aldrich) followed by the reverse-phase HPLC C4 column, as described above. Human ^13^C-^15^N labeled WT α-Syn fibrils were generated, and part of the fibrillar sample was incubated with TNM to induce nitration^69^.

### MAS-ssNMR spectroscopy

MAS-ssNMR experiments were carried out on uniformly ^13^C-^15^N-labeled α-Syn fibrils in both WT and nitrated forms on the 700 MHz Bruker Avance-NEO Spectrometer with a 3.2 mm E-free probe (Bruker) at a temperature of 278 K. DARR experiments were recorded at a spinning rate of 12.5 kHz using a mixing time of 20 ms, and 1 ms of contact time for the cross-polarization (CP) ^1^H-^13^C with RF fields of ν_RF_ = 100 kHz and ν_RF_ = 68.5 kHz for ^13^C and ^1^H, respectively. NCA heteronuclear experiments were acquired at a spinning rate of 12.5 kHz using double CP (DCP) experiments. The ^1^H-^15^N CP conditions were met with ν_RF_ = 45 kHz for ^15^N and ν_RF_ = 57 kHz for ^1^H, and ^15^N-^13^C CP was achieved with a ^15^N RF field of ν_RF_ = 43.75 kHz and ^13^C RF field of ν_RF_ = 18.75 kHz. For all DARR and DCP experiments, decoupling was applied using SPINAL-64 at ν_RF_ = 100 kHz during t_1_ evolution and acquisition, and the complex data were acquired using the States-Time Proportional Phase Incrementation method^69^. NMR data were processed using the NMRpipe software^70^ and analyzed with the Sparky program^71^.

### Animals and surgical procedures

Animal experiments were approved by the State Agency for Nature, Environment, and Consumer Protection in North Rhine Westphalia, Germany. Experiments were conducted in 12-to 14-week old C57BL6/JRJ female mice. Animals were housed in individually ventilated cages in a specific pathogen-free facility and kept on a 12-h light/dark cycle with *ad libitum* access to food and water. For stereotactic injection, mice were anesthetized with isoflurane, and surgery was performed using a stereotactic frame (Stoelting) and a 5 μL Hamilton syringe fitted with a pulled glass capillary tube (outer diameter of 60–80 μm). Animals received a single 2.5 μL injection of human α-Syn PFFs (5 μg), which were injected unilaterally into the right striatum at the following coordinates: 0.2 mm anterior and −2.0 mm lateral to bregma and −2.7 mm ventral relative to the dura. A mouse brain atlas was used as a reference for brain coordinates ^72^. PFFs were injected at a rate of 0.2 μL/min. Following this infusion, the needle was left in position for an additional 5 min before being slowly retracted. The skin incision was then closed with metal clips (Michel clips), and animals were allowed to recover in a heated recovery cage before returning to their home cage. At 12 and 20 weeks postinjection, mice were sacrificed with an intraperitoneal injection of sodium pentobarbital (140 mg/kg) and perfused through the ascending aorta with 4% paraformaldehyde (PFA).

### Tissue preparation and immunohistochemistry

Brains were removed, immersed in 4% PFA for 24 h, and cryopreserved in 30% (w/v) sucrose. Histological staining was carried out on free-floating coronal sections (35 μm). The sections were quenched by incubation in a mixture of 3% hydrogen peroxide and 10% methanol in TBS. Non-specific binding sites were blocked by incubation in 5% normal goat serum for 1 h. Samples were kept overnight at RT in TBS containing 0,25% Triton X, 1% bovine serum albumin (BSA), and primary antibody (see Fig. S1): anti-phosphorylated S129 α-Syn (1:10,000, ab51253; Abcam) or anti-tyrosine hydroxylase (1:1,000, ab152; Merck Millipore). Sections were rinsed with TBS and incubated for 1 h in TBS containing biotinylated secondary antibody (1:200; Vector Laboratories) and 1% BSA. Following treatment with avidin-biotin-horseradish peroxidase complex (ABC Elite Kit; Vector Laboratories), the color reaction was developed using a 3,3′-diaminobenzidine kit (Vector Laboratories). Sections were mounted on coated slides and coverslipped with DPX mounting media (Sigma-Aldrich). Sections used for stereological cell counting were counterstained with cresyl violet.

### Image analysis and stereological cell counting

Brightfield microscopy was used for tissue visualization and histological analyses. These analyses were all performed by investigators blinded to the experimental groups. Phosphorylated α-Syn pathology (pS129-α-Syn) was quantified in the striatum, substantia nigra, and amygdala. Images containing these three brain regions were collected as 20x brightfield stacks using the Zeiss Axio Scan.Z1 microscope (Carl Zeiss). Then analyses were conducted throughout the whole region of interest using every fifth section for a total of 8, 6, and 5 sections for the dorsal striatum, central and basolateral amygdala, and substantia nigra, respectively. In each section, the specific region was delineated using Zen software (Carl Zeiss) and extracted using Adobe Photoshop (CC version 20.0.5). pS129 α-Syn staining was partitioned using Ilastik (version 1.3.0) interactive learning and segmentation tools. The resulting segmented files were thresholded and analyzed using Fiji software (ImageJ version 2.1.0/1.53c).

Unbiased stereology was performed to estimate the number of Nissl-or tyrosine hydroxylase-positive neurons in the substantia nigra. Every fifth section throughout the entire region was analyzed. Delineation of the substantia nigra pars compacta was done using a 4x objective, whereas a 63x Plan-Apo oil objective (numerical aperture = 1.4) was used for neuronal counting. Sections were visualized under the IX2 UCB Olympus microscope, and counts were made using an optical fractionator probe (Stereo Investigator, version 9; MBF Bioscience). The coefficients of error were less than 0.10^73^.

### Preparation and treatment of α-Syn KO and WT hippocampal primary neurons

Primary hippocampal neurons were prepared from WT C57BL/6JRj (Janvier) or α-Syn KO (C57BL/6J OlaHsd; Envigo) pups at postnatal day 0 (P0) and cultured as previously reported^46^. The neurons were seeded in clear black bottom 96-well plates (Falcon), 6-well plates (Falcon) or onto coverslips (CS) (VWR) previously coated with poly-L-lysine 0.1% w/v in water (Brunschwig) at a density of 300,000 cells/mL. After 10 days in culture (DIV 10), α-Syn KO neurons were treated with extracellular human WT or nitrated α-Syn PFFs at a final concentration of 70 nM and their uptake, processing, and clearance were followed from 14 h after PFF addition up to 10 days. In WT neurons, the extracellular human WT or nitrated α-Syn PFFs were added 6 days after plating the neurons (DIV 6) at a final concentration of 70 nM as previously described^32, 46, 47, 74^. PBS was used as a negative control in α-Syn PFFs experiments. All procedures were approved by the Swiss Federal Veterinary Office (Authorization Nos. VD 3392 and VD3694).

### Subcellular fractionation and WB analysis of α-Syn KO and WT hippocampal primary neurons

PBS-or PFFs-treated α-Syn KO and WT neurons were lysed in 1% Triton X-100/TBS supplemented with protease inhibitor cocktail, 1 mM PMSF, and phosphatase inhibitor cocktails 2 and 3 (Sigma-Aldrich). After sonication (0.5 s pulse at 20% amplitude, 10 times), cell lysates were incubated on ice for 30 min and centrifuged at 100,000 *g* for 30 min at 4°C. The supernatant (soluble fraction) was collected while the pellet was washed in 1% Triton X-100/TBS, sonicated for 5 s pulse at 20% amplitude four times, and centrifuged for another 30 min at 100,000 *g*. The supernatant was discarded, whereas the pellet (insoluble fraction) was resuspended in 2% SDS/TBS supplemented with protease inhibitor cocktail, 1 mM PMSF, and phosphatase inhibitor cocktails 2 and 3, and sonicated (0.5 s pulse at 20% amplitude, 15 times). The Bicinchoninic acid protein assay was performed to quantify the protein concentration in the soluble and insoluble fractions before the addition of Laemmli buffer 4x. Protein samples were separated on 1-mm-thick 16% Tricine gels (Life Technologies) for 2 h at 120 V. The proteins were transferred to 0.2 µm nitrocellulose membranes (Amersham) using a semidry transfer system (Bio-Rad) under constant current (0.5 A) and a voltage of 25 V. The membranes were blocked for 1 h at RT in Odyssey blocking buffer (Li-COR Biosciences) and probed with the respective primary antibodies overnight at 4°C (see Supplemental Fig. 1 for all the information related to the antibodies used in our study). After three washes with PBS containing 0.1% (V/V) Tween 20 (Fluka), the membranes were incubated with secondary goat anti-mouse or anti-rabbit antibodies conjugated to Alexa fluor 680 or 800 (1:20,000; Li-COR Biosciences). Then the membranes were washed three times with PBS containing 0.01% (V/V) Tween 20, and scanned on a LiCOR scanner (Li-COR Biosciences). The levels of total α-Syn or pS129 α-Syn were detected by WB, quantified using Image Studio software (RRID:SCR_015795; Li-COR Biosciences), and normalized to the relative protein levels of actin. All experiments were independently repeated three times.

### ICC

After α-Syn PFFs treatment, primary hippocampal neurons were washed twice with PBS, fixed in 4% PFA for 20 min at RT, and then immunostained as previously described^46^. The seeded aggregates were detected using mouse monoclonal pS129 (81A; Biolegend) or rabbit monoclonal pS129 (MJF-R13; Abcam) antibodies. Neurons were counterstained with MAP2 antibody, and the nucleus was stained with DAPI. The cells plated on CS were examined with a confocal laser scanning microscope (LSM 700; Carl Zeiss Microscopy) with a 40x objective and analyzed using Zen software (RRID:SCR_013672). The cells plated in the clear black bottom 96-well plates were imaged using the Nikon 10×/ 0.45, Plan Apo, CFI/60 of the IN Cell Analyzer 2200 (GE Healthcare), a high-throughput imaging system equipped with a high-resolution 16-bits sCMOS camera (2048×2048 pixels)^32, 46^. For each independent experiment, images of three wells were acquired per tested condition, and in each well, nine fields of view were imaged. Each experiment was repeated at least three times independently. Quantification of the pS129 intensity formed in PFFs-treated WT hippocampal neurons was performed as previously described^46^. In brief, images were analyzed using Cell profiler 3.0.0 software (RRID:SCR_007358) for quantifying the level of pS129-positive seeded aggregated formed in MAP2-positive cells. The pipeline of this analysis has been fully detailed in Mahul-Mellier et al. (2020)^46^.

### Statistical analyses

Data from at least three independent experiments were used for all statistical analyses. Statistical analyses on normally distributed data were performed using *t*-tests or one-way analysis of variance (ANOVA) followed by the Tukey’s Honest Significant Difference posthoc test or using two-way ANOVA with Tukey’s multiple comparisons for comparisons between two or more groups (*in vivo* data). P < 0.05 was considered statistically significant.

## Supporting information

Supplemental informations

## Acknowledgments

We thank the EPFL Interdisciplinary Center for Electron Microscopy CIME facility for the use of their electron microscopy facility. We also would like to thank the Proteomics Core Facility (PTP, EPFL) for access to the MALDI-TOF instrument. The authors thank Laura Demmer for assistance with *in vivo* tissue preparation and Lorène Aeschbach (LMNN, EPFL) and Yllza Jasiqi (LMNN, EPFL) for their help with the preparation of the primary neuronal cultures used in this work. We also thank Galina Limorenko (LMNN, EPFL) for her help in creating the Graphical abstract (Figure 5).

## Author contributions

Conceptualization: HAL

Experimental design: HAL, SD, ALMM, DDM, ADS, and SOS

Data curation: HAL, SD, SOS, ALMM, AU, GF, STK, AC, JB, MB, IR, ADS, DDM

Formal analysis: HAL, SD, SOS, ALMM, AU, GF, STK, AC, JB, MB, IR, ADS, DDM

Validation: HAL, SD, SOS, ALMM, AU, GF, STK, AC, JB, MB, IR, ADS, DDM

Visualization: HAL, SD, SOS, ALMM, AU, GF, STK, ADS, DDM

Supervision: HAL

Writing—original draft: HAL and SD

Writing—review & editing: HAL, SD, JB, SOS, ALMM, AU, GF, STK, IR, ADS, DDM

Funding acquisition: HAL

Project administration: HAL

Resources: HAL

## Data and material availability

All data are available in the main text or supplementary materials.

