## Supplemental informations for "Post-fibrillization nitration of alpha-synuclein abolishes its seeding activity and pathology formation in primary neurons and *in vivo*"

**Running title:**

Effect of post-fibrillization nitration on α-Synuclein

**Keywords:** alpha‐synuclein (α‐synuclein), Parkinson's disease, post‐translational modifications (PTMs), post-fibrillization nitration, fibrils.

**Legends**

**Figure S1**

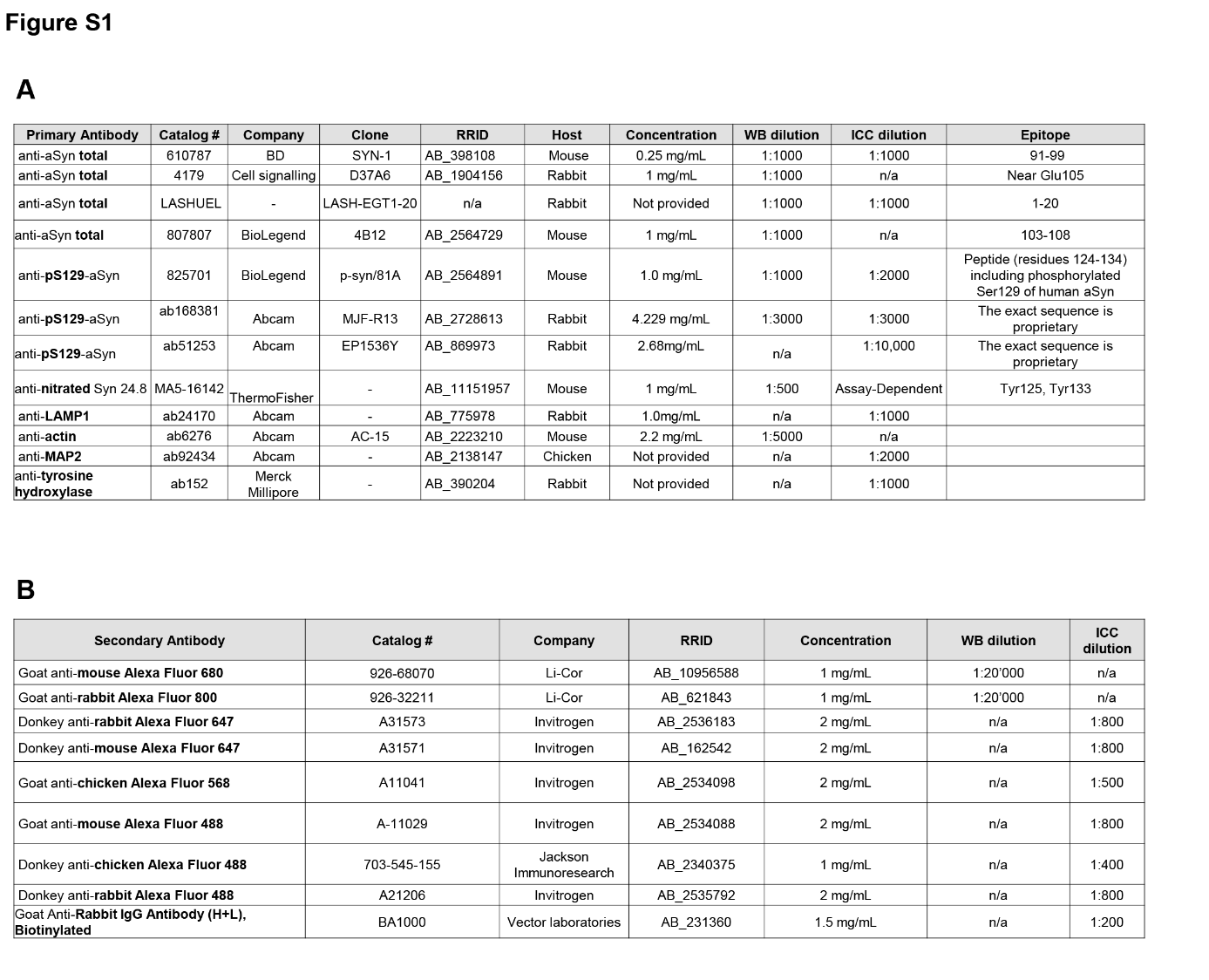

**Supplemental Figure 1: List of the primary and secondary antibodies used in this study.**

Primary antibodies (**A**) and secondary antibodies (**B**) used for WB or IHC and ICC analyses.

**Figure S2**

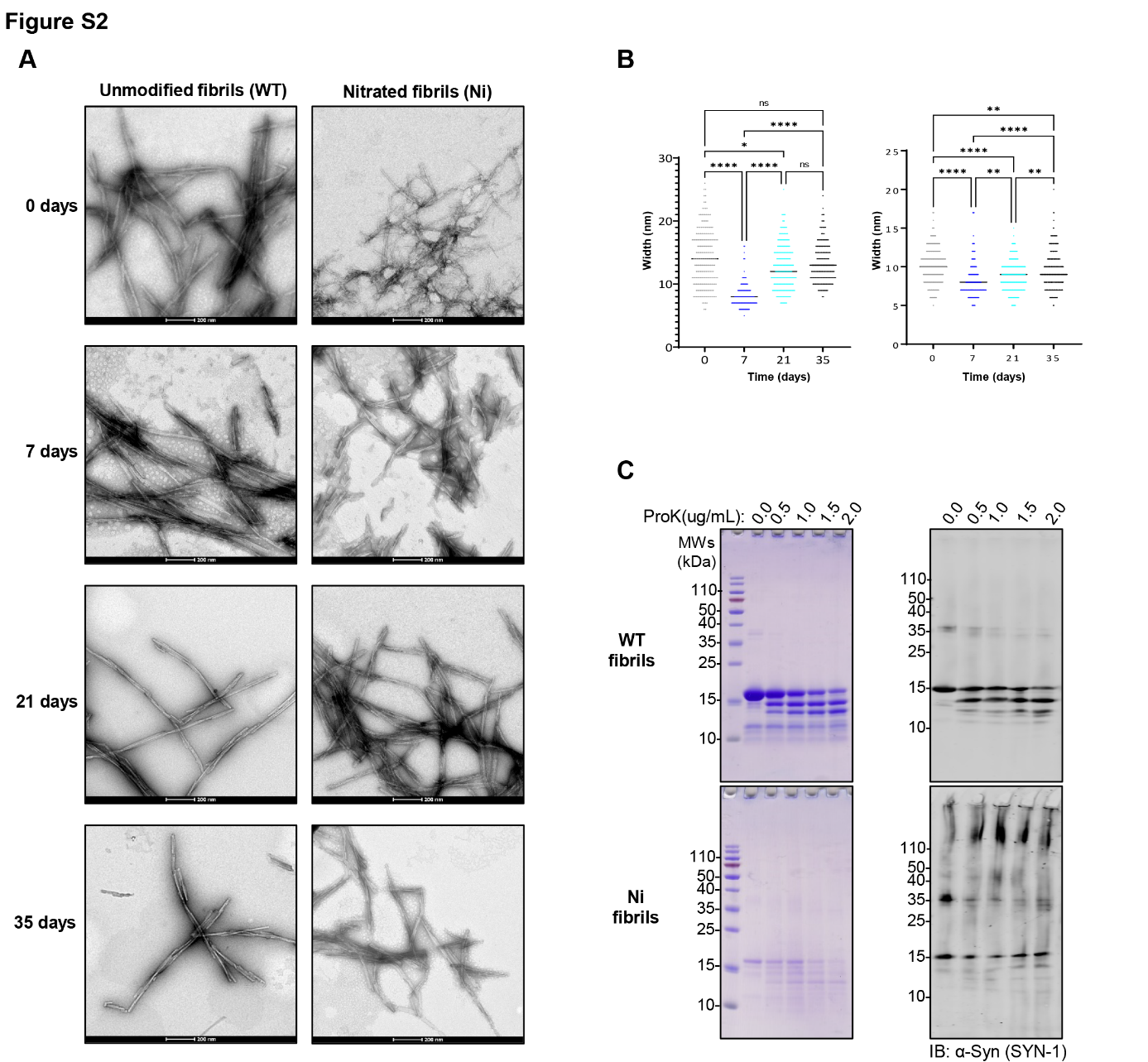

**Supplemental Figure 2: Post-fibrillization nitration does not alter the structure of α-Syn PFFs.**

**A-B.** TEM images (**A**) and quantification of the width (**B**) of unmodified and nitrated fibrils at different time points. **C.** Coomassie stainings and Western Immunoblot analysis of α-Syn fibrils (10 ug/mL) treated with proteinase K (PK) at increasing concentration for 30 min. Scales bars = 200 nm.

**Figure S3**

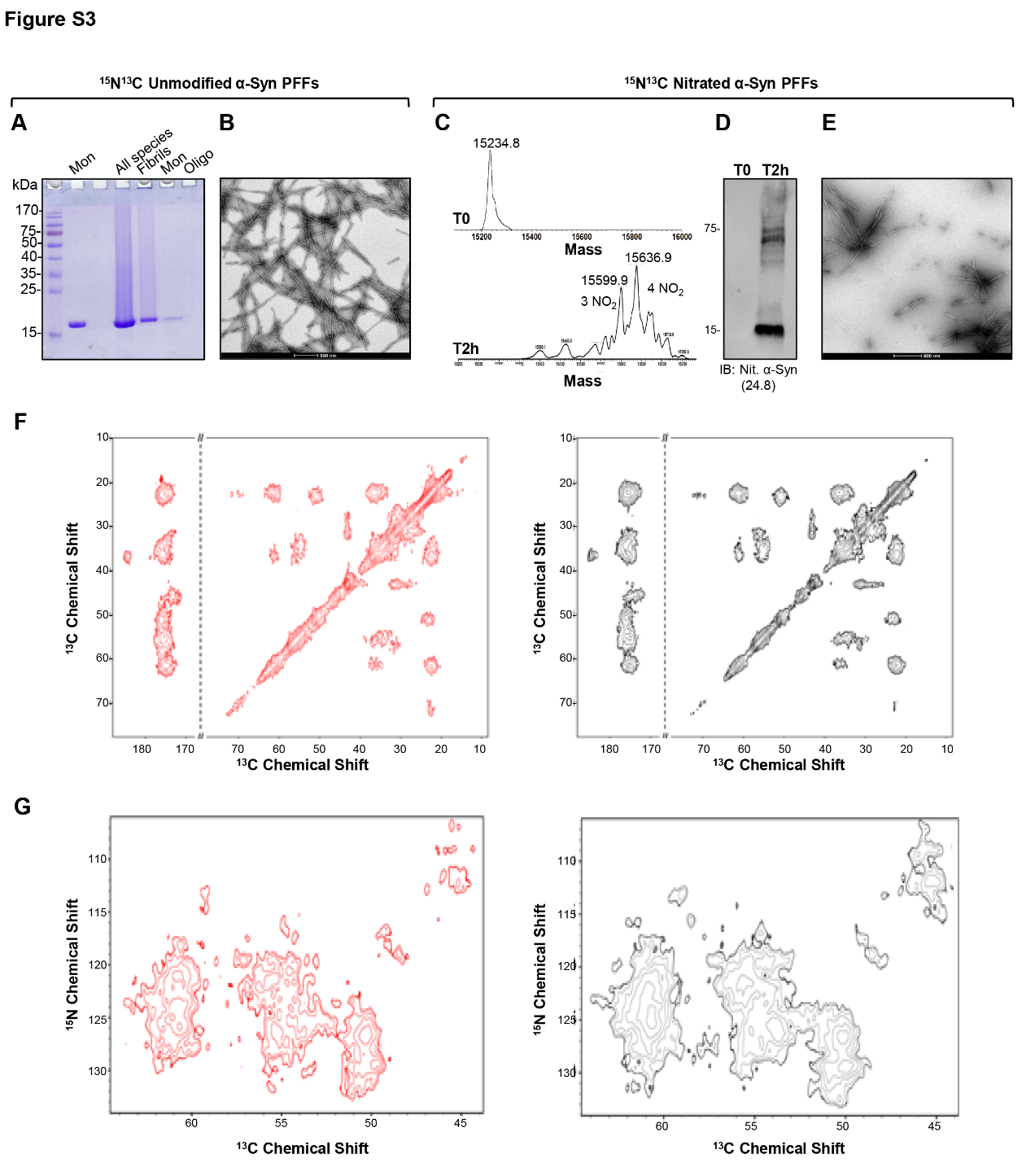
**Supplemental Figure 3:** **Characterization of the human** **^15^N^13^C unmodified and nitrated PFFs by in vitro NMR studies.** **A.** Coomassie staining of ^15^N^13^C unmodified monomeric α-Syn (first line, before fibrillization), all species present in the solution after sonication (second line), isolated PFFs (third line), and monomers (fourth line), or oligomers (fifth line) released after sonication. **B.** TEM image of ^15^N^13^C unmodified α-Syn PFFs. **C.** ESI-LC/MS analysis of the extent of TNM-mediated nitration of ^15^N^13^C α-Syn PFFs after 2h of incubation. ^15^N^13^C α-Syn PFFs were boiled at 95°C for 10 minutes, spun down by centrifugation (max speed, 4°C, 10 min), and the supernatant was injected into the ESI-MS for analyses. **D.** WB analysis on unmodified (T=0) and nitrated ^15^N^13^C α-Syn PFFs (T=2h). α-Syn nitration antibody 24.8 was used to detect post-fibrillization nitration on PFFs. **E.** TEM image of ^15^N^13^C nitrated α-Syn PFFs. Scale bars = 200 nm. The images are qualitative. **F.** Individual ^13^C-^13^C DARR ssNMR spectra of unmodified (red) and nitrated (black) are reported in Figure 1G. **G.** Individual ^13^C-^15^N NCA ssNMR spectra of unmodified (red) and nitrated (black) are reported in Figure 1H.

**Figure S4 – part I**

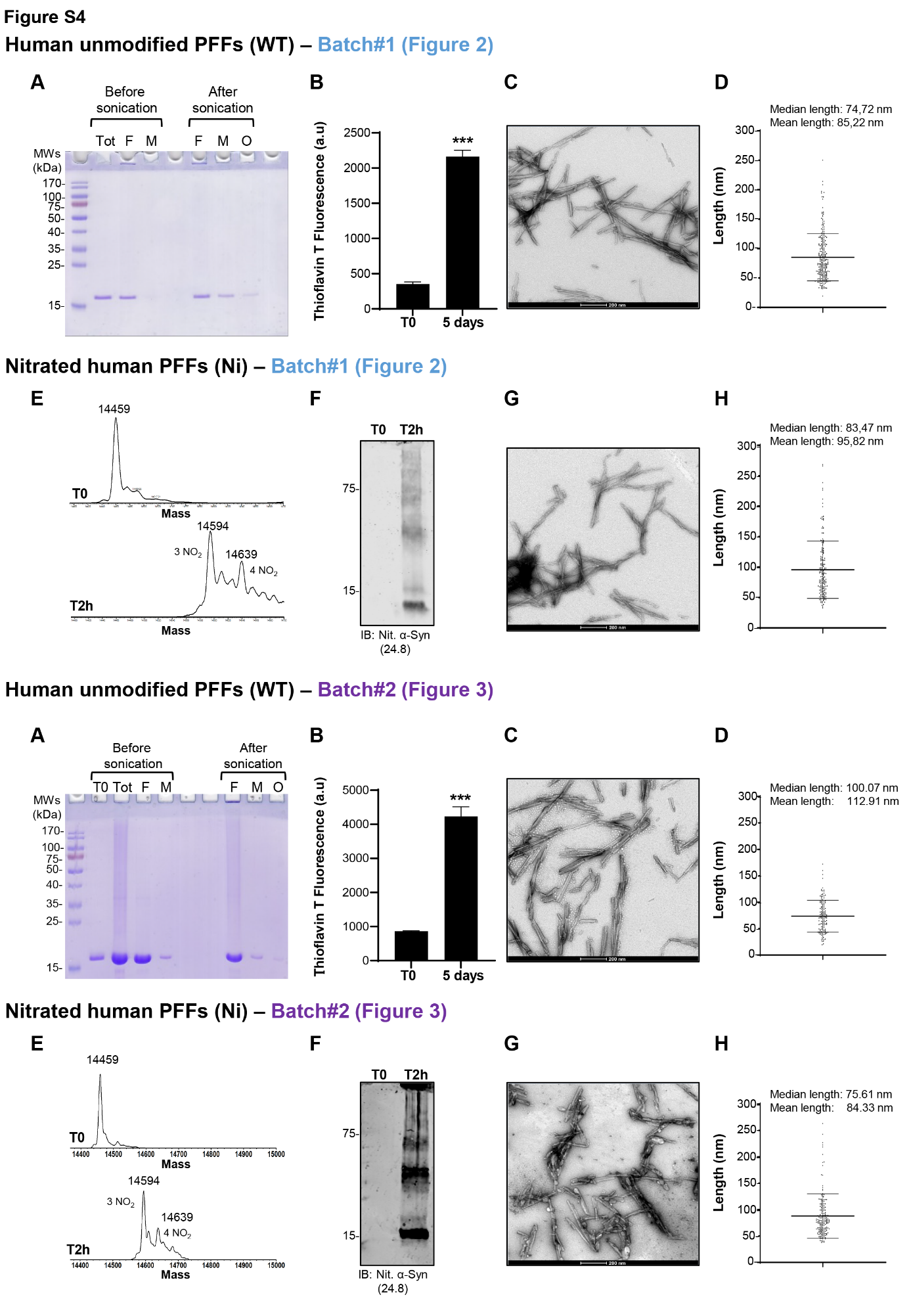

**Figure S4 – part II**

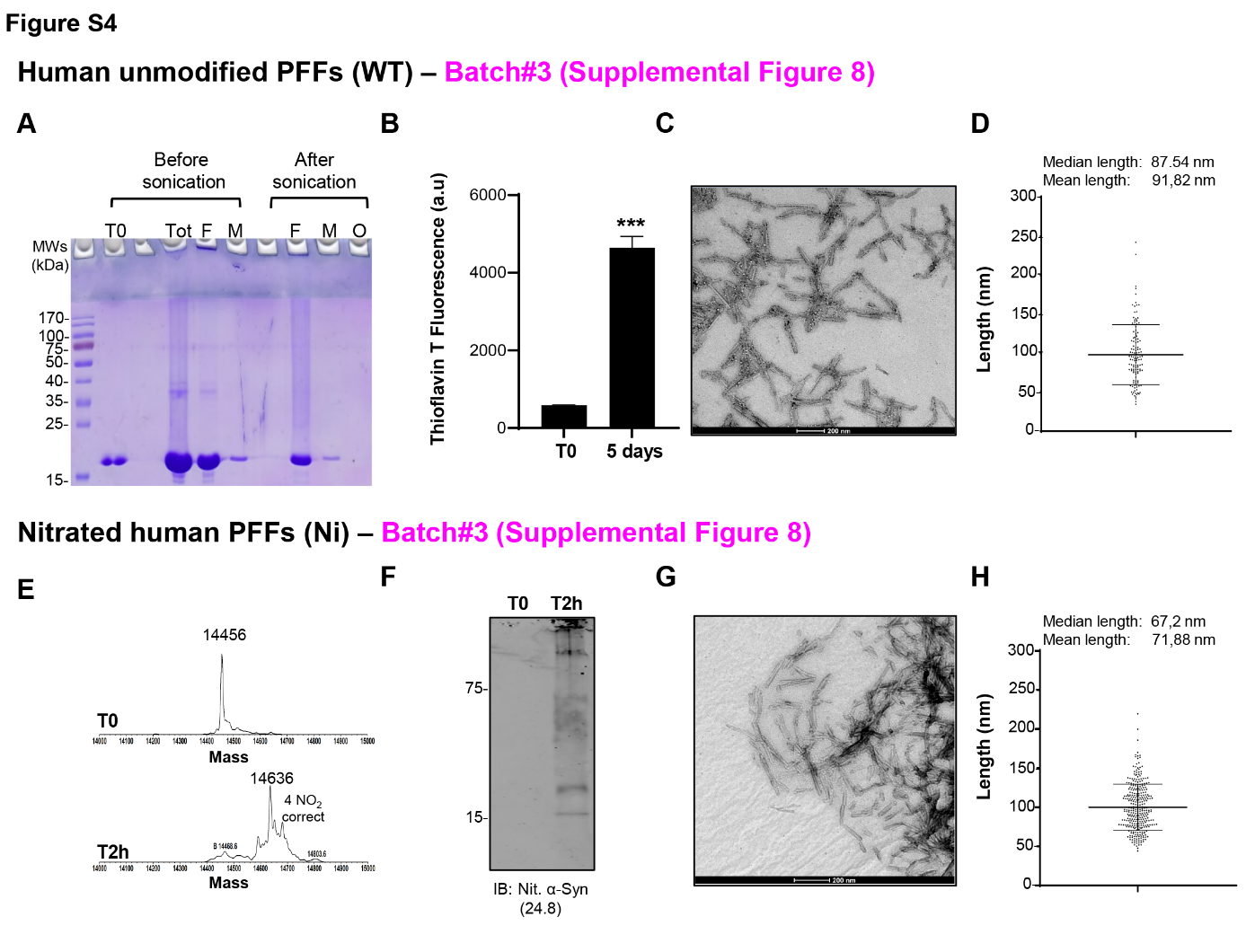

**Supplemental Figure 4: Preparation and characterization of unmodified or nitrated α-Syn PFFs used in the in vivo and cellular studies.**

Three independent batches of unmodified or nitrated human WT PFFs were prepared (batch #1 for the in vivo studies and batches #2 and #3 for the cellular studies) and characterized as follows. **A.** Unmodified monomeric α-Syn was incubated for 5 days at 37°C under constant agitation at 1000 rpm. The resulting fibrils were then sonicated, and the amount of released monomers and oligomers after sonication was quantified by SDS-PAGE and Coomassie blue staining as previously described (Kumar, Donzelli et al. 2020). **B.** Fibril formation was also assessed by ThT fluorescence-based aggregation assay. All data represent the average ± SD (n=3). Statistical significance was determined using a one-way ANOVA test followed by Tukey HSD posthoc test. **C, G.** The structure of the fibrils unmodified (**C**) or nitrated (**G**) after sonication was monitored by TEM. Scale bars = 200 nm. **D, H**. The length of the sonicated fibrils unmodified (**D**) or nitrated (**H**) was quantified for the TEM images. **E-F.** Nitration of the WT α-Syn fibrils was confirmed by ESI-LC/MS and WB analyses. **E.** ESI-LC/MS analysis of the extent of TNM-mediated nitration of α-Syn fibrils after 2h of incubation. α-Syn fibrils were boiled at 95°C for 10 minutes, spun down by centrifugation (max speed, 4°C, 10 min), and the supernatant was injected into the ESI-MS for analyses. **F.** WB analysis on unmodified (T=0) and nitrated fibrils (T2h). The α-Syn nitration antibody 24.8 was used to detect post-fibrillization nitration on fibrils.

**Figure S5**

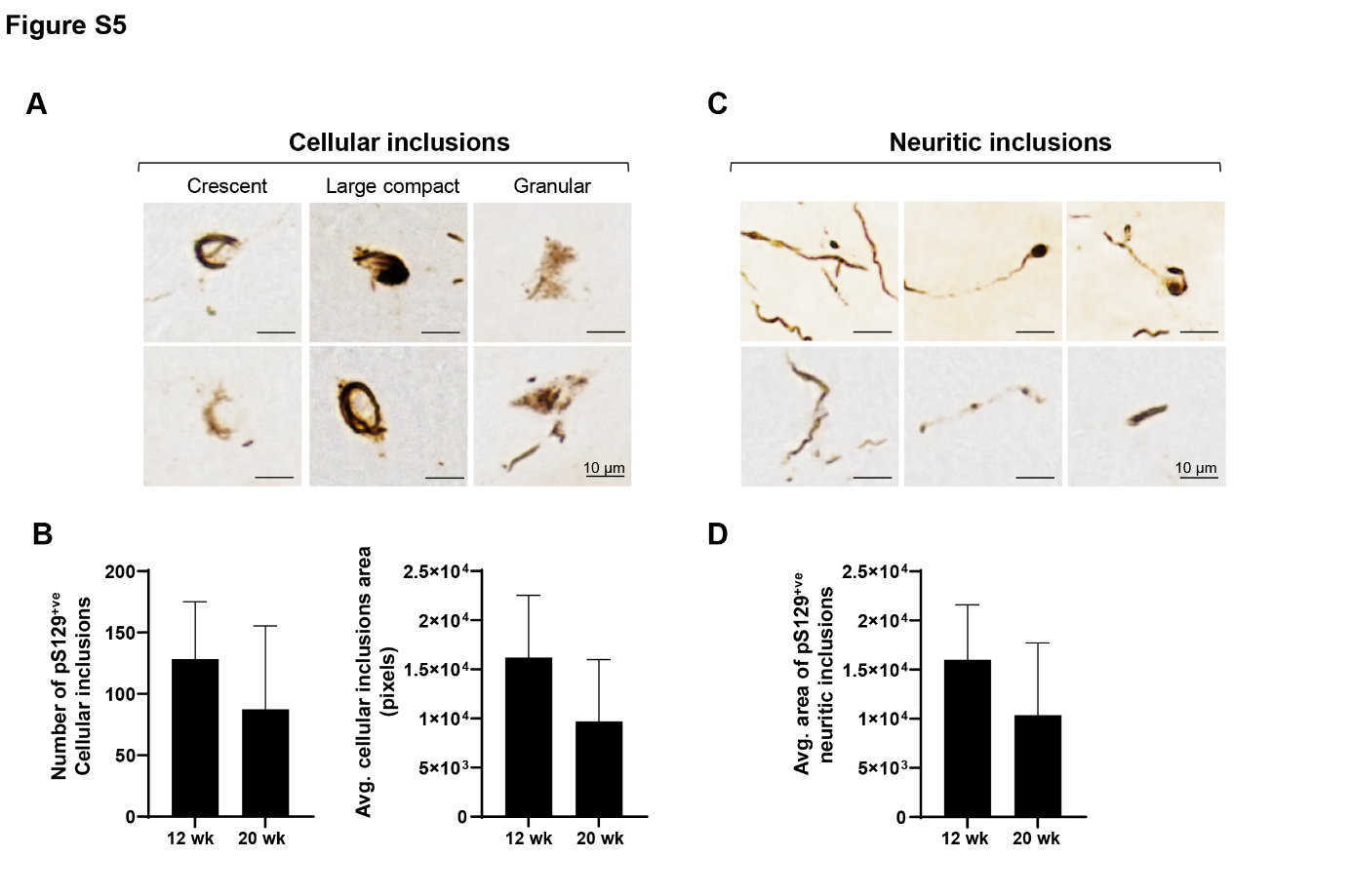

**Supplemental Figure 5: Characterization of unmodified PFFs in vivo.**

Unmodified human PFFs were injected unilaterally into the mouse striatum, and animals were sacrificed at 12 or 20 weeks post-injection (n = 8-9 animals/time point). Midbrain sections were stained with an anti-pS129 antibody (EP1536Y). **A.** Representative images show neuronal cell bodies in the substantia nigra pars compacta containing inclusions that were different in shape and immunoreactivity. **B.** The total number of pS129-α-Syn-positive cellular inclusions and the total area occupied by these inclusions were measured in equally spaced midbrain sections encompassing the entire substantia nigra pars compacta. **C.** Representative images show neuritic inclusions in the substantia nigra pars compacta. **D.** The total area occupied by nigral neuritic inclusions was measured. Scale bars = 10 µm. Graphs show means +/- SD.

**Figure S6**

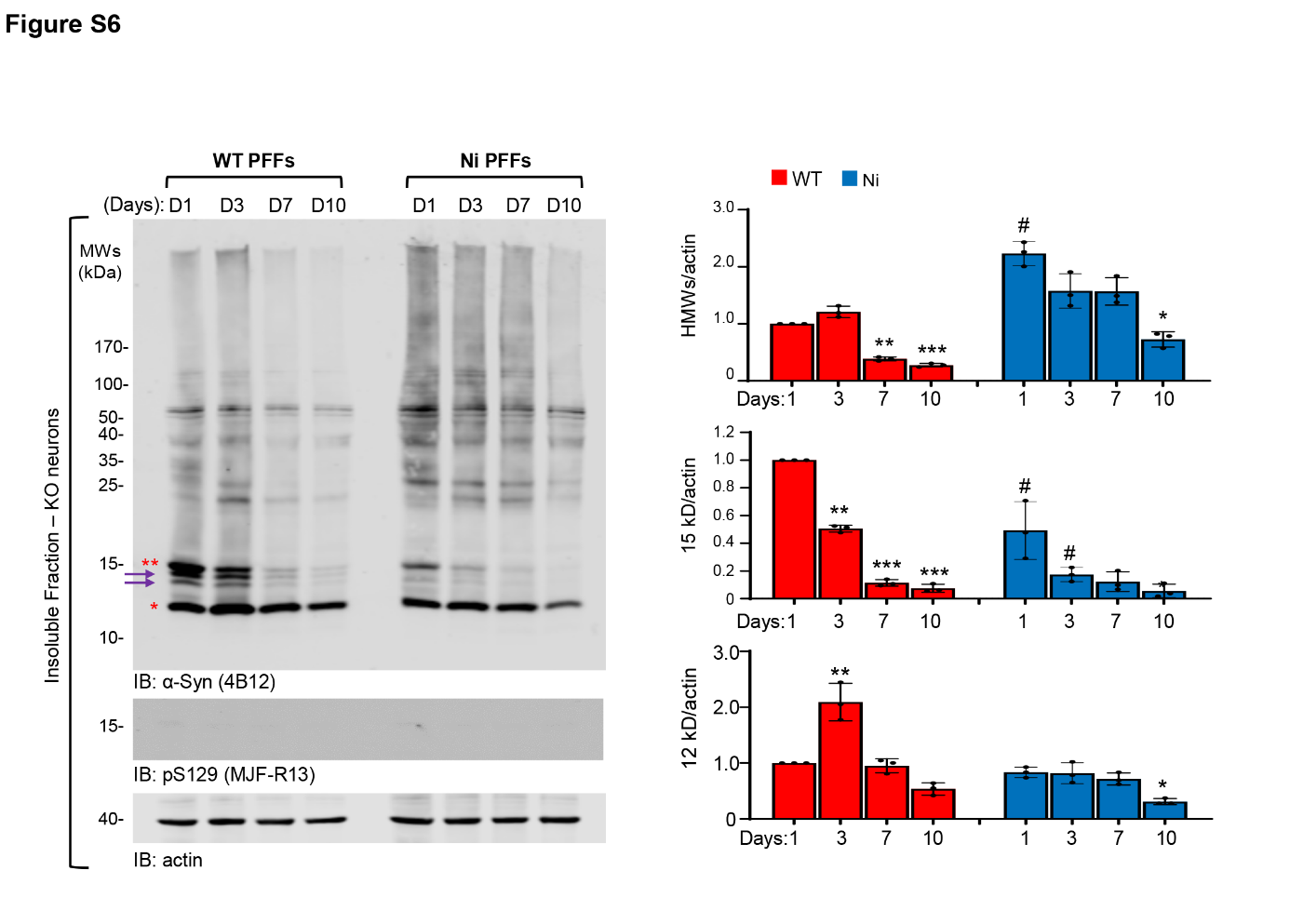

**Supplemental Figure 6: Post-fibrillization nitration does not interfere with the uptake of α-Syn PFFs, their processing (C-terminal cleavage) or the general patterns of their clearance after internalization.**

aSyn KO primary hippocampal neurons at 10 days in vitro (DIV 10) were treated with the indicated PFFs (70 nM) or PBS for up to 10 days. WB analysis shows that nitration of the human WT PFFs does not alter their internalization, their C-terminal cleavage to ~12 kDa fragment, or their clearance overtime in primary neurons. The quantification of the immunoblots using the total α-Syn antibody 4B12 shows no significant difference between the WT or the nitrated WT PFFs. Results are the mean ±SD of biological replicates (n=3). Statistical significance was determined using a one-way ANOVA test followed by Tukey HSD posthoc test.

**Figure S7**

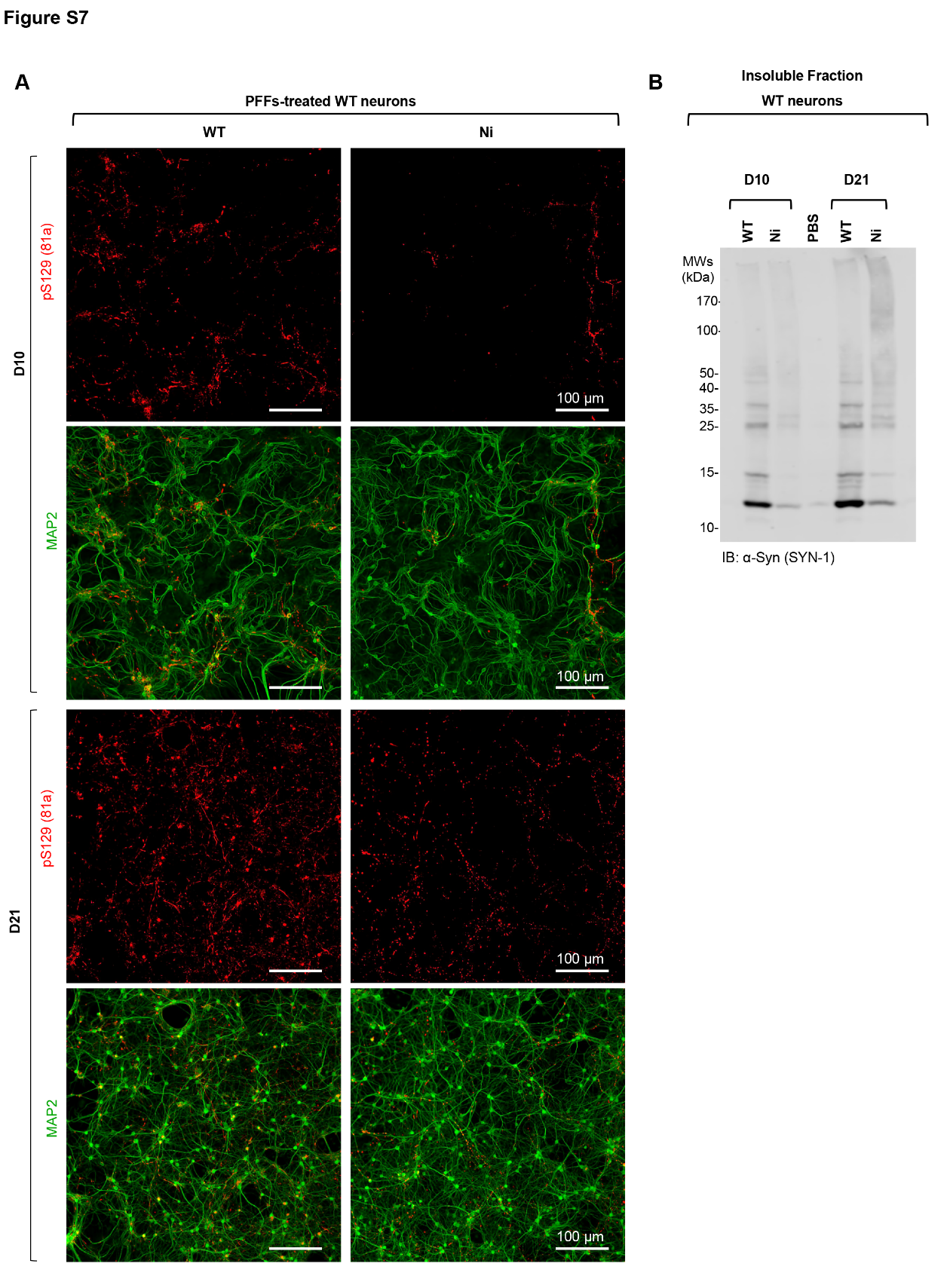

**Supplemental Figure 7: Post-fibrillization nitration reduces the seeding capacity of α-Syn PFFs in primary neurons.**

WT hippocampal primary neurons were treated with 70 nM of WT or Ni α-Syn PFFs (batch #1) for 10 (D10) or 21 days (D21). The seeding capacity of the PFFs was evaluated by ICC (**A**) or by WB analyses (**B**). **A.** Representative images of the seeded aggregates formed in the MAP2-positive neurons. ICC was performed using the pS129 antibody (81A) and imaged by the IN Cell Analyzer 2200. Scale bars = 100 μm. **B.** WB analysis of the insoluble fractions of WT neurons treated with WT or Ni human PFFs. The level of seeded aggregates was detected with an antibody specific for total α-Syn (SYN-1).

**Figure S8**

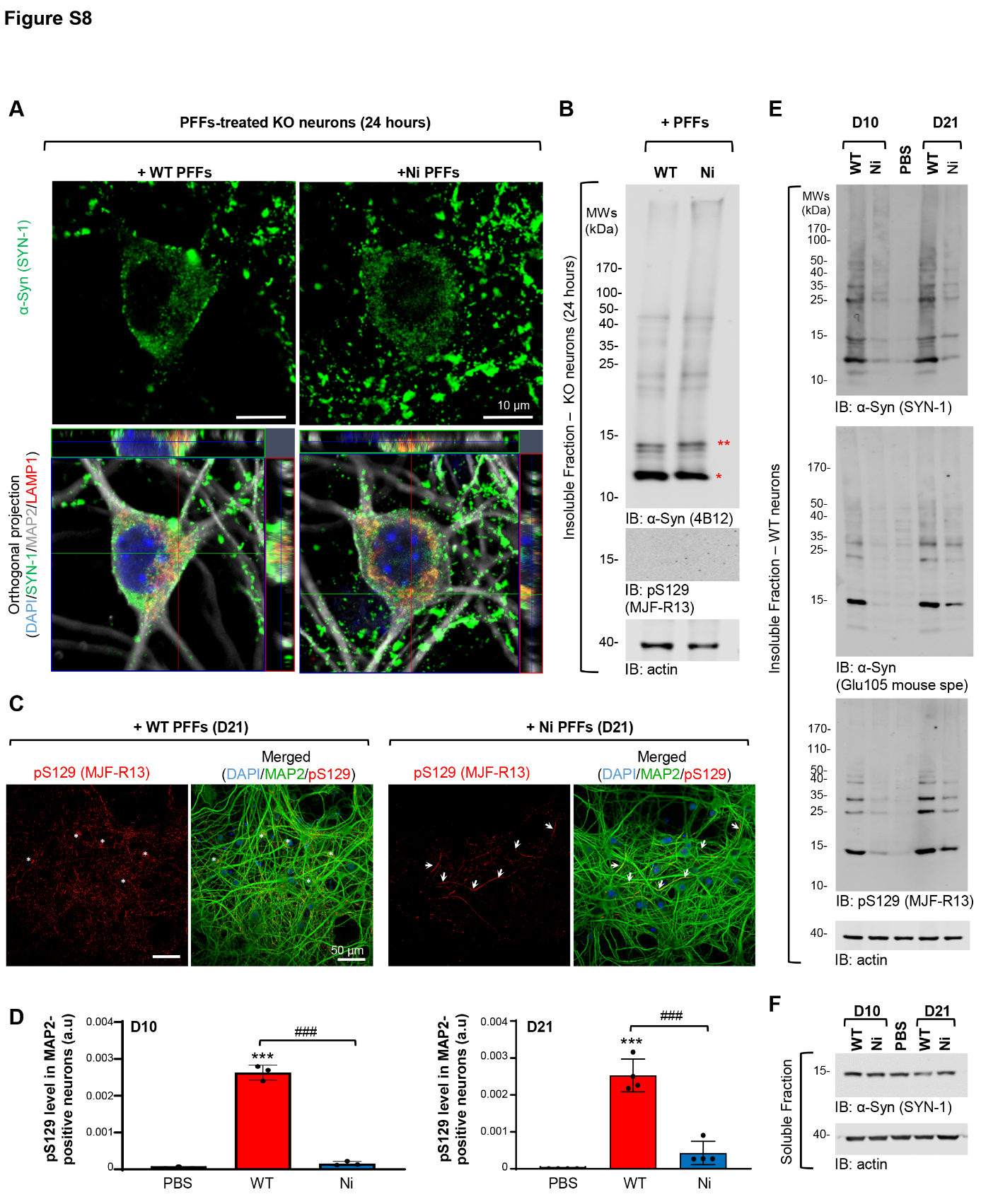

**Supplemental Figure 8: Uptake, processing and seeding capacity of the WT and nitrated PFFs in primary hippocampal neurons.**

α-Syn KO (**A-B**) or WT (**C-G**) hippocampal primary neurons were treated with 70 nM of WT or Ni α-Syn PFFs (batch #3, see characterization in Supplemental Figure 4) for 24 hours (D1, **A-B**) and up to 21 days (D21, **C-E**). Control neurons were treated with PBS. The uptake, processing and seeding capacity of the PFFs were evaluated by ICC (**A**), High content imaging (HCA, **C-D**) and WB (**B, E-F**) analyses. **A.** ICC analysis shows the internalization of WT and Ni α-Syn PFFs into the MAP2-positive α-Syn KO neurons. The PFFs were detected by total α-Syn (SYN-1) antibody, the neurons were counterstained with microtubule-associated protein (MAP2) antibody, and the nuclei with DAPI. The Late endosome/lysosomes were stained with LAMP1 antibody. Scale bars = 10 μm. **B.** WB analysis of the insoluble fractions of α-Syn KO neurons treated with WT and Ni α-Syn PFFs for 24 hours shows the detection of the 12 kDa (one red star), 15 kDa (double red stars), and HMWs bands by the total α-Syn antibody (4B12). **C.** The seeded aggregates formed in the cell bodies (white star) and in the neurites (white arrows) of the MAP2-positive neurons were detected by ICC using the pS129 antibody (81A) and imaged by the IN Cell Analyzer 2200. Scale bars = 50 μm. **D.** The level of pS129- α-Syn-positive seeded aggregates in (**C**) was quantified by HCA. For each independent experiment (n=3), a minimum of two wells was acquired per condition, and nine fields of view were imaged per well. **E.** WB analysis of the insoluble fractions of WT neurons treated with WT or Ni human PFFs. The level of seeded aggregates was detected with an antibody specific for mouse α-Syn (Glu105) or with a pS129 antibody (MJF-R13). SYN-1 was used to stain total α-Syn. **F.** WB analysis of the soluble fractions of WT neurons treated with WT or Ni PFFs shows the decrease of the endogenous α-Syn (15 kDa), specifically in the WT PFFs-treated neurons at D21. The levels of α-Syn were normalized to the actin.

**Figure S9**

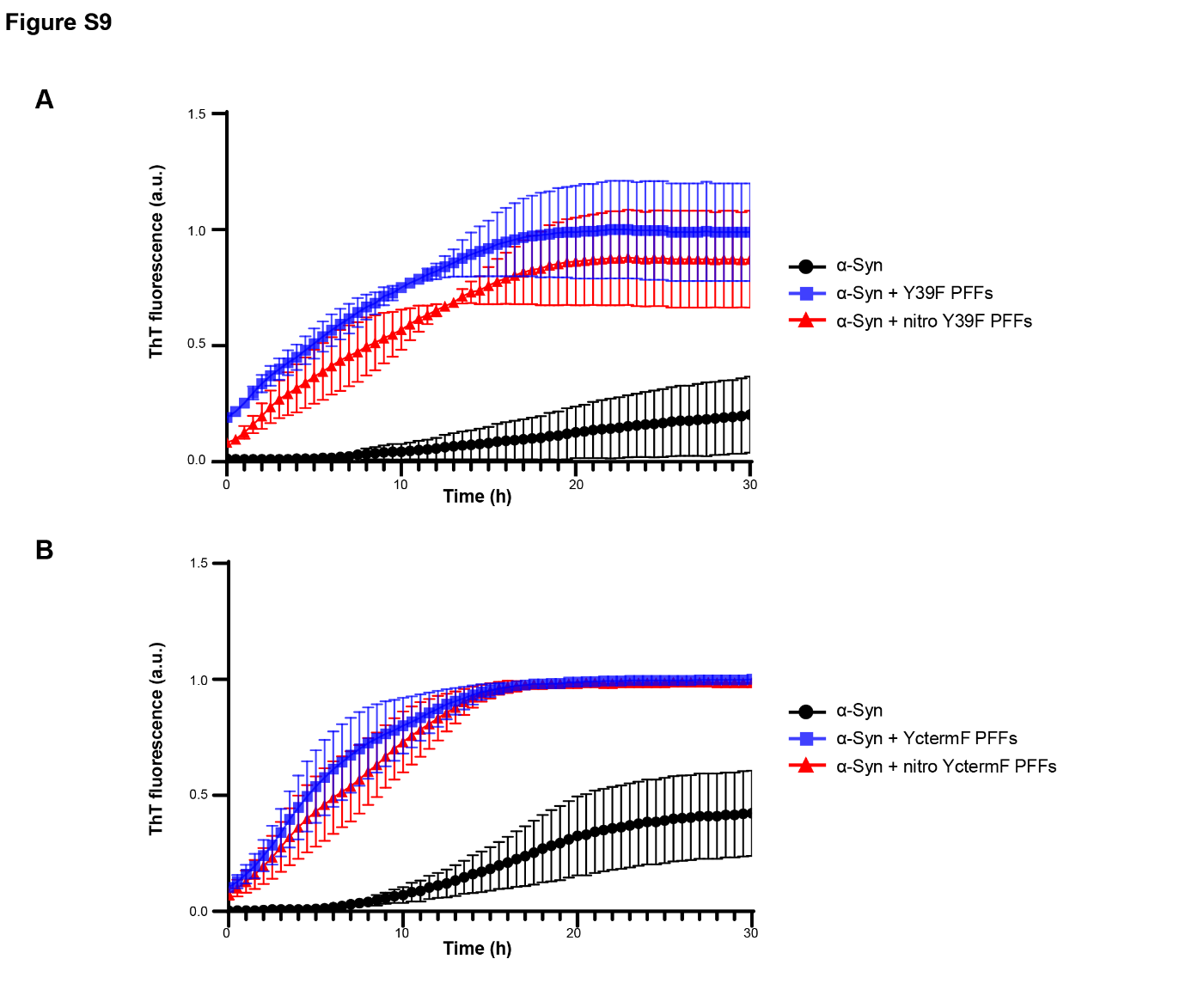
**Supplemental Figure 9: Seeding capacity of α-Syn fibrils specifically nitrated at YC-term or at Y39.**

**A.** Fluorescent Thioflavin T (ThT) assay of recombinant monomeric α-Syn, (black line), monomeric α-Syn + Y39F fibrils (blu line), and monomeric α-Syn + nitrated Y39F fibrils (red line). **B.** Fluorescent Thioflavin T (ThT) assay of recombinant monomeric α-Syn, (black line), monomeric α-Syn + YctermF fibrils (blu line), and monomeric α-Syn + nitrated YctermF fibrils (red line).

**Figure S10**

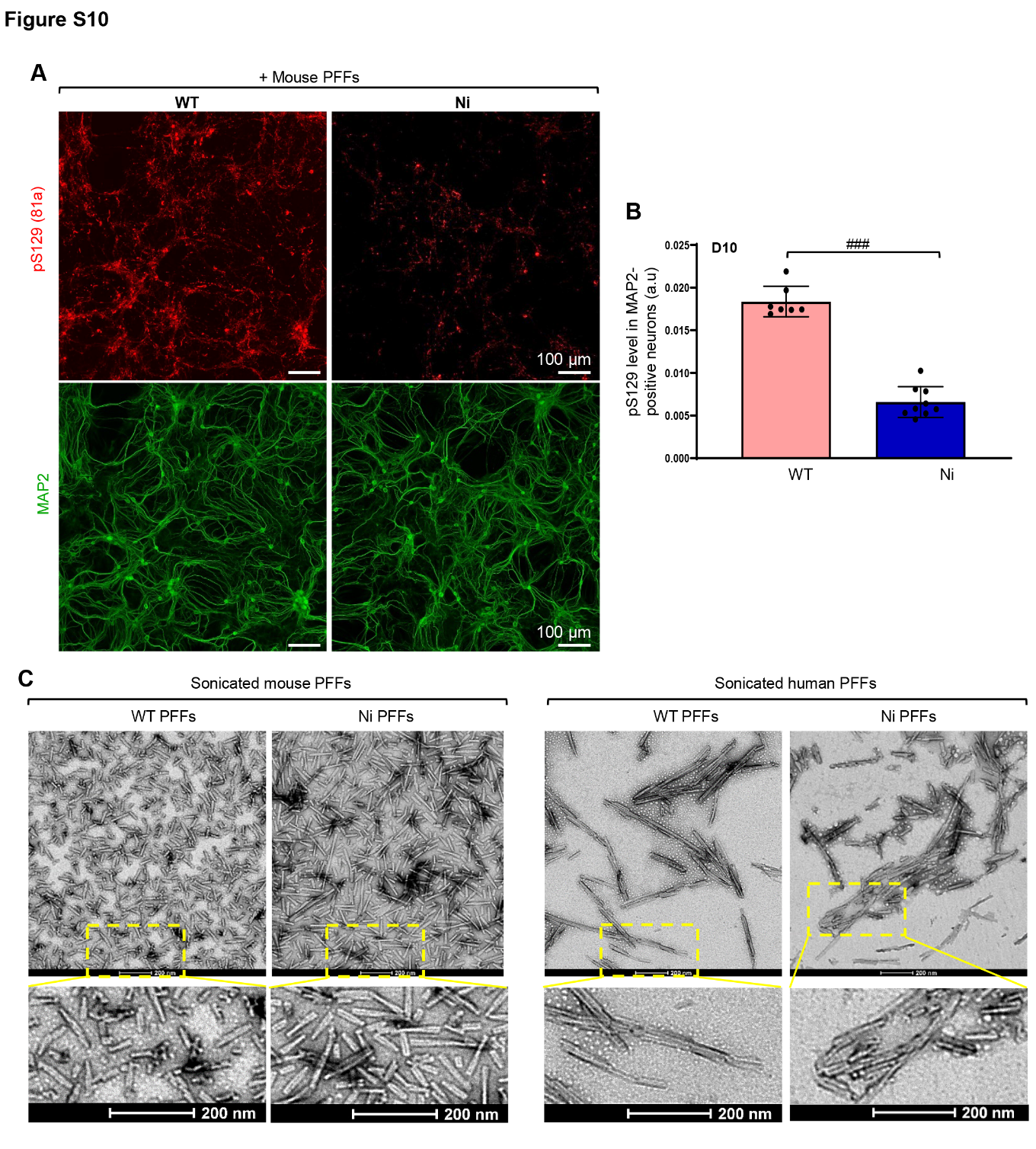
**Supplemental Figure 10: Seeding capacity of unmodified or nitrated α-Syn mouse fibrils.**

WT hippocampal primary neurons were treated with 70 nM of mouse WT or Nitrated (Ni) α-Syn PFFs. **A-B.** The seeding capacity of the PFFs was evaluated by ICC (**A**) and High content imaging quantification (**B**). **A.** The seeded aggregates formed in the WT or Ni mouse PFFs-treated MAP2-positive neurons were detected by ICC using the pS129 antibody (81A) and imaged by the IN Cell Analyzer 2200. Scale bars = 100 μm. **B.** The level of pS129-α-Syn-positive seeded aggregates in (**A**) was quantified by HCA. For each independent experiment (n=3), a minimum of two wells was acquired per condition, and nine fields of view were imaged per well. The graphs represent the mean +/- SD of a minimum of 3 independent experiments. **B.** ###p<0.0001 (one-way ANOVA followed by Tukey HSD posthoc test, WT vs. Ni PFFs-treated neurons). C) TEM representative images of mouse and human sonicated PFFs that have been left unmodified (WT) or that have been nitrated (Ni). Scale bars = 200 nm.
